# The phage protein paratox is a multifunctional metabolic regulator of *Streptococcus*

**DOI:** 10.1101/2024.08.02.605684

**Authors:** Tasneem Hassan Muna, Nicole R. Rutbeek, Julia Horne, Ying Wen Lao, Oleg V. Krokhin, Gerd Prehna

## Abstract

*Streptococcus pyogenes*, or Group A Streptococcus (GAS), is a commensal bacteria and human pathogen. Central to GAS pathogenesis is the presence of prophage encoded virulence genes. The conserved phage gene for the protein paratox (Prx) is genetically linked to virulence genes, but the reason for this linkage is unknown. Prx inhibits GAS quorum sensing and natural competence by binding the transcription factor ComR. However, inhibiting ComR does not explain the virulence gene linkage. To address this, we took a mass-spectrometry approach to search for other Prx interaction partners. The data demonstrates that Prx binds numerous DNA binding proteins and transcriptional regulators. We show binding of Prx *in vitro* with the GAS protein EndoS1 (SpyM3_0890) and the phage protein JM3 (SpyM3_1246). An EndoS1:Prx complex X-ray crystal structure reveals that EndoS1 and ComR possess a conserved Prx binding helix. Computational modelling predicts that the Prx binding helix is present in several, but not all, binding partners. Namely, JM3 lacks the Prx binding helix. As Prx is conformationally dynamic, this suggests partner-dependent binding modes. Overall, Prx acts as a metabolic regulator of GAS to maintain the phage genome. As such, Prx maybe a direct contributor to the pathogenic conversion of GAS.

## INTRODUCTION

*Streptococcus pyogenes* (Group A Streptococcus or GAS) is a Gram-positive bacteria that is both a human commensal and pathogen^1–3^. The bacteria colonize the throat and skin in 1 to 5% of healthy adults and up to 20% of children^2,4^. GAS causes pharyngitis (strep throat), toxic shock syndrome, rheumatic fever, scarlet fever, and is the leading cause of flesh-eating disease world-wide^3,5^. Overall GAS is responsible for more than 600 million pathogenic infections each year, with more than 500,000 deaths due to invasive disease^5,6^. However, as GAS is a human-restricted, or human-only, bacteria^7^ this raises questions as to both why and how the bacteria leaves a commensal niche. The mechanisms for the transition from a commensal into a pathogen are highly studied questions for GAS, as the decision to destroy your host is thought to be an evolutionary disadvantage^2,8^.

GAS virulence is in large part due to bacteriophages that can enter and then later leave the bacterial genome^9,10^. GAS phages are temperate double-stranded DNA phages belonging to the *Siphoviridae* family^9,10^. These phages can switch between a lysogenic cycle where they insert their genome into the GAS host genome (prophage) or enter a lytic cycle where they create new phage particles and ultimately spread to new hosts. Furthermore, the GAS phage contains genes that encode toxins that are essential to GAS pathogenesis^7,11^. These include superantigens such as SpeA (streptococcal pyrogenic exotoxin) that result in toxic shock syndrome^7^, and other virulence factors such as phospholipases and DNases^3,11,12^. Once infected by a GAS phage, the bacterial cell becomes more virulent and has the potential to convert into a pathogen. Due to this, infection by phages is thought to be a mechanism for both the rise of new pathogenic GAS strains and the primary driving force for GAS clonal diversity^9,13^.

Related to the evolution of GAS through phage infection is another prominent question in GAS biology. Specifically, why GAS are resistant to other mechanisms of genetic manipulation^14^. GAS exhibit resistance to both artificially induced genetic competence such as electrocompetence, and natural genetic competence in a laboratory setting. This is despite GAS possessing all the genes required for the uptake and processing of environmental DNA^15–17^. However, both questions are in part answered by the activity of the small phage protein named paratox (Prx).

In GAS, the ComRS quorum sensing system regulates natural competence or uptake of environmental DNA^15,16^. ComR is a receptor and transcription factor that is a member of the Rgg subgroup of the RRNPP protein family^18^, and ComS is a peptide pheromone signal that is processed into a mature form termed XIP (sig<u>X</u> inducing <u>p</u>eptide)^15,16^. At the molecular level, ComR undergoes an extensive conformational change upon binding XIP^19,20^. Binding of XIP to ComR releases its DNA-binding-domain (DBD) which allows ComR to dimerize and recognize DNA.^20^ ComR is then able to induce the expression of ComS (XIP) creating a positive feedback loop. Activated ComR is also able to induce expression of the alternative sigma-factor SigX, which in turn upregulates expression of the GAS competence machinery^16^. Prx binds the ComR DBD directly with nano-molar affinity blocking interaction of ComR with DNA^21,22^. Additionally, Prx is able to bind and inhibit ComR regardless of if ComR is in the inactive monomer conformation or the active XIP-bound dimer conformation^22^. Furthermore, Prx expression is also induced by SigX which creates a negative feedback loop to counter positive feedback loop of the GAS quorum sensing signal^16,21^. As ComRS regulates the transcription of genes for the DNA uptake and recombination machinery, Prx inhibits both GAS quorum sensing and GAS natural competence^21^.

In addition to inhibiting natural competence, deletion of *prx* has been observed to increase GAS electrocompetence by ∼100,000-fold^21^. This suggests that Prx may alter GAS membrane permeability (DNA uptake) and/or regulate DNA processing pathways other than ComRS. However, unlike the role of Prx in natural competence the molecular mechanism for the electrocompetence phenotype of Prx is yet to be described. Also unanswered is why *prx* is genetically linked to prophage toxin genes such as *speA*^21,23^. The gene *prx* is encoded adjacent to a GAS toxin gene at the 3’ end of the prophage and the two genes are in “linkage disequilibrium.” Specifically, *prx* and its toxin gene partner remain together as one genetic cassette during homologous recombination^23^. As such, *prx* and the toxin gene are never separated in the creation of new infectious phage particles during the lytic cycle. Given that Prx inhibits DNA uptake, we had previously hypothesized that the biological role of Prx maybe to protect the integrity of the linked toxin gene from damage or loss by recombination^21,22^. However, our current biochemical data do not provide a solid explanation for why *prx* is explicitly linked to GAS virulence genes.

It is not uncommon for bacteriophage proteins to have more than one biological or biochemical function. For example, the phage protein Aqs1 from *Pseudomonas aeruginosa* inhibits both quorum sensing and pilus assembly^24^. Considering this and a lack of data explaining the *prx*-toxin linkage and the *prx* deletion electrocompetence phenotype, we asked if Prx has other binding partners in addition to ComR. To search for Prx interaction partners we performed pull-downs of purified Prx with GAS lysates. Mass-spectrometry analysis of the co-precipitated peptides showed that Prx may interact with a multitude of proteins. These include several putative DNA binding proteins, many of which contain a helix-turn-helix (HTH) DBD domain. We then show direct binding of Prx to two DNA-binding proteins, one from the GAS host (EndoS1 or SpyM3_0890) and a second encoded by the prophage (JM3 or SpyM3_1246). Next, we obtained a crystal structure of EndoS1 and a co-crystal structure of an EndoS1:Prx complex which together demonstrate how Prx inhibits EndoS1 from binding DNA. Additionally, a structural analysis of known experimental Prx co-crystal complexes aided by AlphaFold complex predictions, reveals a conserved ɑ-helix structural motif for Prx-binding that is found in several GAS proteins. However, JM3 lacks the ɑ-helix binding motif suggesting that Prx may have other mechanisms of partner recognition. Overall, our work demonstrates that Prx influences the biochemistry of both the GAS host and the bacteriophage by blocking the interaction of gene-regulatory and DNA processing proteins with DNA. Given this and the direct linkage of *prx* to toxin genes, the metabolic manipulation of GAS by Prx likely contributes directly to the transition of *Streptococcus pyogenes* from a commensal into a pathogen.

## RESULTS

### Prx interacts with several GAS and phage DNA binding proteins

In a previous study we observed that deletion of *prx* increased GAS electrocompetence by approximately 100,000-fold^21^. This phenotype combined with the observation that phage proteins often have multiple biological functions led us to hypothesize that Prx may have additional binding partners. To test this hypothesis, we searched for potential Prx interaction partners by pull-down assays with different GAS lysates. Our experiments used strain MGAS315 as our Prx recombinant protein expression construct was cloned from the same strain^21^. Purified Prx-6His was immobilized on nickel-affinity resin as bait with empty beads used as controls. Lysates were created separately from GAS cells stimulated with XIP to both induce Prx expression and natural competence gene expression^16,21^, and from GAS cells stimulated with mitomycin C to induce phage exit from the genome (lytic cycle)^9,10^. We also performed a pull-down with the spent media from GAS cells stimulated by mitomycin C to capture proteins lost by phage-induced cell lysis. Proteins that co-precipitated with immobilized Prx were eluted from beads, digested, and detected by mass-spectrometry. The results of each experiment are presented in Figure S1, S2, and S3 with a curated protein list of initial interest displayed in Figure 1. Results are listed by log expectation value (more negative, more probable)^25^, with a cut-off value of ‘-10’ set for potential hits given the large data set. The initial Prx binding partner hits displayed in Figure 1 and Figures S1-S3 were selected by relation to known Prx function (DNA-binding)^21^ and phage-biology, and detection levels in +Prx experiments as compared to -Prx experiments. Ribosomal proteins were filtered due to their common presence as false positives pull-down assays^26^. Additionally, the top two hits based on expectation value of all three pull-downs are included (SpyM3_0271 and SpyM3_0575). As shown in Figure 1 and Figures S1-3, the pull-down data for the several dozen potential Prx interaction partners are all very high-confidence, especially as most hits have no detected peptides in the bead-only controls (-Prx). Furthermore, in these experiments Prx was observed to pull-down its previously known binding partner ComR^21^, but only from mitomycin C induced lysates (Figure 1, orange bar). Prx also co-precipitated with other paratox proteins. Importantly, MGAS315 has 6 *prx* genes and the three listed Prx hits are all different genes from our recombinant Prx-6His construct. Additionally, as we have shown that Prx does not oligomerize *in vitro*^21^, this suggests that there is either an *in vivo* condition within GAS that allows Prx to recognize itself or that the co-precipitation is indirect. Namely, that the other Prx proteins are bound to the same protein target as Prx. This is a distinct possibility as DNA-binding proteins often dimerize for interaction with DNA which could provide two Prx binding sites^20,27,28^.

**Figure 1:**
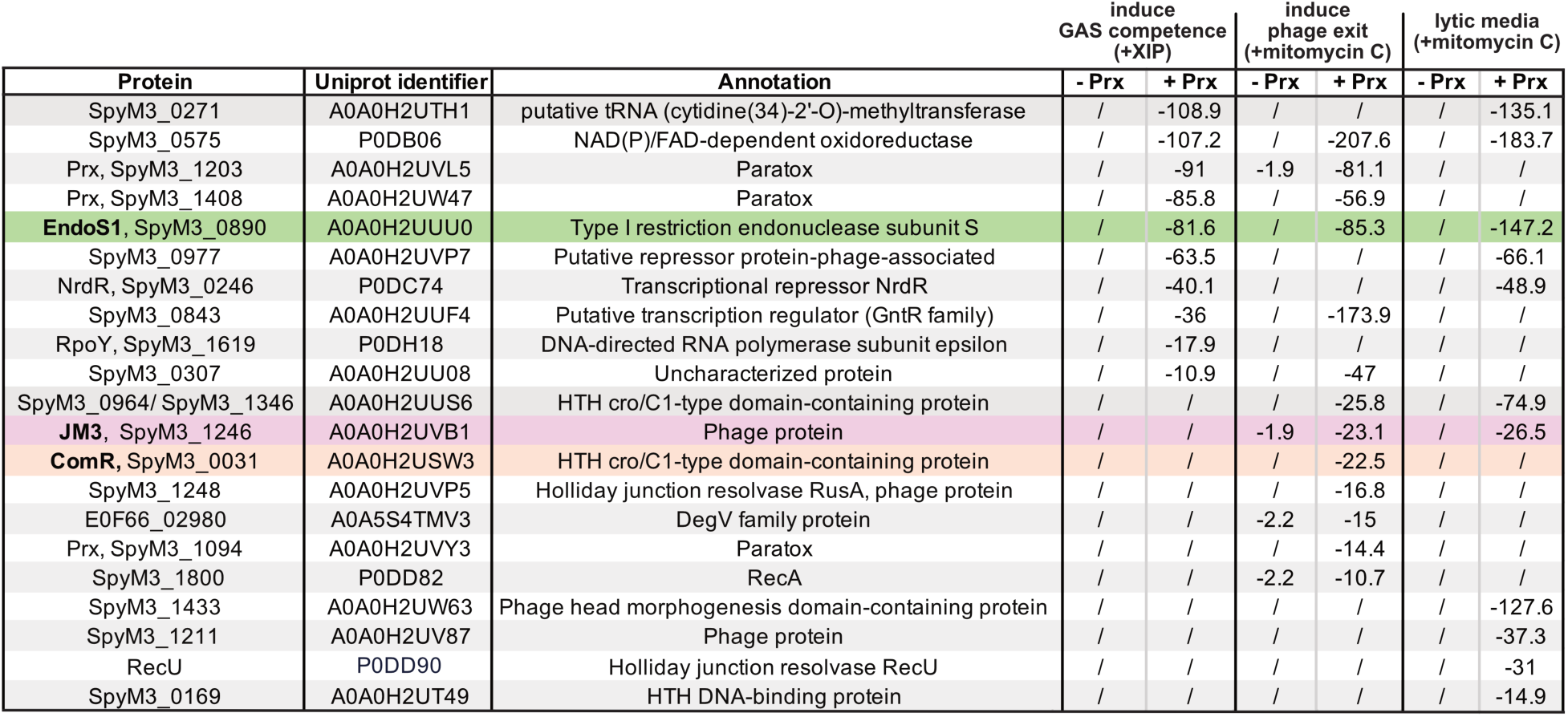
Prx binding partners identified from *Streptococcus pyogenes.* Purified Prx bound to nickel-agarose beads (+Prx) was used to co-precipitate proteins from GAS extracts. Extracts were stimulated with either XIP to induce GAS natural competence, or mitomycin C to induce phage protein expression and lytic exit. The media from cells growing in mitomycin C was also used to co-preciptate proteins (lytic media). Empty beads (-Prx) was used as a control. Protein hits were identified by mass-spectrometry and are listed by protein name, Uniprot identifier, and putative annotations. Data is reported as the Log expectation value, where a larger negative value indicates a higher confidence protein hit. No peptides detected by mass-spectrometry is indicated by a backslash. The data is this table is a curated list of the complete results in Figures S1, S2, and S3.

In Figure 1, the overall pattern appears to be that Prx co-precipitates with a large number of DNA and RNA binding proteins, with each protein being involved in distinct biochemical pathways. These include proteins such as SpyM3_0977, NrdR^27^, and SpyM3_0843 that are possible transcriptional regulators, and proteins like SpyM3_1248 and RecU involved in DNA recombination^28^. Additionally, the detected proteins span both GAS proteins and phage proteins such as a phage head morphogenesis protein (SpyM3_1433). Morphogenesis proteins are known to assist in phage assembly^29^. The top hit (SpyM3_0271) is a putative tRNA methyltransferase suggesting that Prx may also interact with RNA binding proteins. Prx also co-precipitated with several other tRNA methyltransferases including TrmR from XIP induced lysates (E0F66_03415, Figure S1), SpyM3_0634 from mitomycin C induced lysates (Figure S2), and SpyM3_1500 from the mitomycin C or phage induced lysate media (Figure S3). This of course assumes that SpyM3_0271 and the other methyltransferases are not non-specific bead-proteome false positives^26^. It is also important to state that all of the listed putative Prx binding partners in Figure 1 are annotated by predicted function bioinformatically, and to our knowledge the majority have not been experimentally characterized.

Additional Prx binding patterns are also readily apparent from Figure 1. For example, some protein hits such as SpyM3_0890 are observed in both GAS metabolic states. Namely, induced natural competence (XIP) and mitomycin C simulated phage exit (Figure 1, green). However, other pull-down hits such as SpyM3_1246 were identified only when the phage has been induced to by mitomycin C (Figure 1, purple). Specifically, there are many putative Prx binding partners that are only found when GAS cells have been induced to produce phage proteins (see both +mitomycin C experiments, Figure 1 and S1-3). Another apparent pattern is that many of the Prx binding partners contain helix-turn-helix (HTH) DNA-binding domains (DBD). This includes SpyM3_0964/1346 (which has the same specific HTH subclass as ComR, cro/C1-type), SpyM3_0169, SpyM3_0977, SpyM3_0843, and SpyM3_1036 (Figure1 and Figure S1). For the last three proteins, the HTH is annotated in the Uniprot entry and readily observable in the AlphaFold model. Of note, SpyM3_0964 and SpyM3_1346 have same amino acid sequence but are encoded as distinct genes making them impossible to differentiate by this experiment. As HTHs are a hallmark of DNA binding and are typically diagnostic of transcription factors^18–20^, the pull-down data strongly suggest that Prx may exhibit wide-spread metabolic control over GAS.

### Prx directly binds the *Streptococcus* protein EndoS1 and the phage protein JM3

The pull-down experiments in Figure 1 and Figures S1-3 represent only a probability of protein binding, albeit a very high interaction confidence. Additionally, as the pull-downs were performed with GAS lysates many of the putative Prx binding partners maybe indirect through a third binding partner. In fact, this possibility is highlighted by the observation that other Prx proteins were detected. As such, the next step required was a validation of the listed hits as direct Prx binding proteins. For validation, both SpyM3_0890 and SpyM3_1246 were selected. For clarity, SpyM3_0890 was renamed to EndoS1 (<u>Endo</u>nuclease <u>S</u>-subunit-like-1) given its predicted annotation (Figure 1) and SpyM3_1246 will be referred to a JM3 (<u>J</u>ulia and <u>M</u>una construct 3). EndoS1 was selected as it is a GAS protein present in both XIP and mitomycin C stimulated lysates, and JM3 was chosen as it was found only from phage induced lysates (Figure 1). Additionally, although both proteins are predicted to bind DNA only JM3 contains an HTH DBD domain similar to ComR^19,22^. Furthermore, EndoS1 was of specific interest given its functional annotation. The S-subunit (<u>S</u>pecificity-subunit) of type I restriction endonucleases recognizes DNA and then recruits the methylation (M) and restriction (R) subunits to enzymatically modify the target DNA^30–32^. Typically, the function of a fully assembled RMS, or restriction-modification enzyme, is to degrade foreign DNA and to help protect a bacteria from phage infection^31^. Finally, both proteins were also of interest given their genomic locations. EndoS1 is encoded near several metabolite transporters and membrane permeases, and JM3 is encoded within the phage recombination machinery near another Prx binding partner, RusA (SpyM3_1248) (Figure S4). As such, EndoS1 and JM3 may be linked to the known *prx* deletion phenotype of DNA processing and membrane permeability related to increased electrocompetence^21^.

To test if Prx binds EndoS1 and JM3 directly, both *endoS1* and *JM3* were amplified from MGAS315 genomic DNA and subcloned into recombinant protein expression vectors. EndoS1 was created with a C-terminal 6His-tag for metal affinity purification and JM3 was made as a fusion with a protease cleavable N-terminal GST. In our hands JM3 was unstable as a 6His construct. After purification and removal of GST from JM3, each protein was incubated with a 1.5x molar excess of Prx and binding assayed by size exclusion chromatography (SEC). As shown in Figure 2A, EndoS1 (23 kDa) is readily purifiable and forms a complex with Prx (8 kDa). This is evident in the SEC traces that show a clear increase in size and loss of an EndoS1 peak (blue), indicative of stable complex formation with Prx (green). Additionally, the accompanying SDS-PAGEs gels of the SEC elution fractions show a definite EndoS1:Prx complex. Similarly, JM3 (10 kDa) was also readily purifiable and forms a stable complex with Prx. Analogous to EndoS1:Prx, the JM3:Prx complex is clearly observed by a shift to a larger species in solution by SEC (dark purple). The JM3:Prx complex was also verified by SDS-page gel (Figure 2B). These data show that in addition to the *S. pyogenes* competence regulator ComR, Prx also forms complexes with the GAS protein EndoS1 and the phage protein JM3.

**Figure 2:**
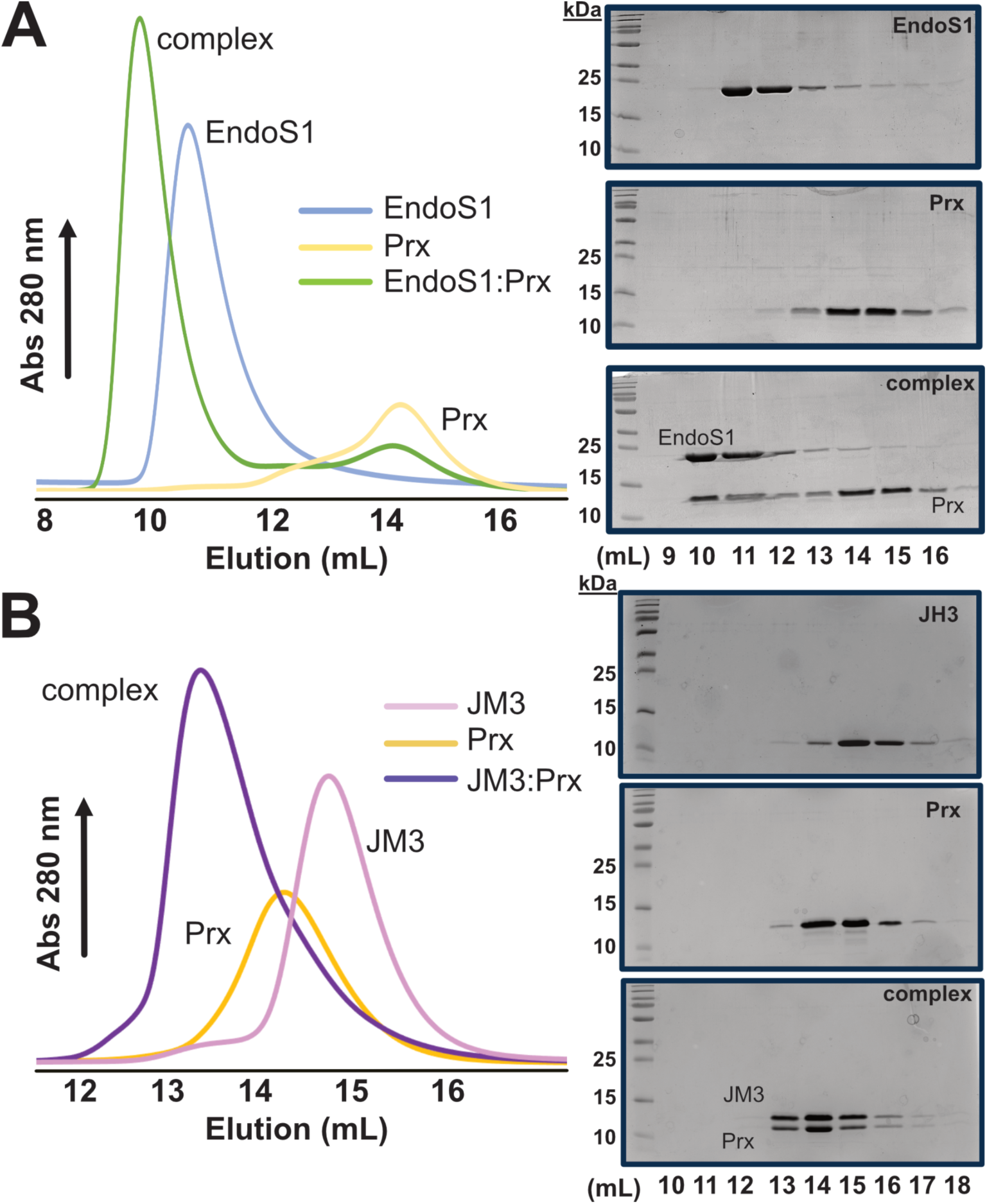
Prx binding assays with purified EndoS1 and JM3. **(A)** Size-exclusion chromatography (SEC) binding assay of EndoS1 with Prx. EndoS1 (blue) mixed with a 1.5 molar excess of Prx (gold) produces a clear shift in elution volume indicative of complex formation (green). The adjacent panels show Coomassie stained SDS-PAGE gels corresponding to each SEC trace. **(B)** SEC binding assay of JM3 with Prx. JM3 (pink) mixed with a 1.5 molar excess of Prx (gold) produces a shift in elution volume indicative of complex formation (purple). The adjacent panel shows Coomassie stained SDS-PAGE gels corresponding to each SEC trace. For all SDS-PAGE gels, SEC fractions collected from the respective elution volumes in mL are shown.

### EndoS1 is structurally distinct from type I restriction endonuclease S-subunits

To further characterize the newly discovered Prx binding partner EndoS1, we determined an X-ray crystal structure of EndoS1 (Figure 3 and Table 1). EndoS1 crystallized as a dimer in the asymmetric unit, with each monomer consisting of an N-terminal globular head and a C-terminal dimerization helix (Figure 3A). In the crystal structure, the EndoS1 dimer appears to be formed exclusively through knob-in-hole hydrophobic packing interactions of the dimerization helices (Figure 3B). Given this, the observed dimeric structure was verified by size-exclusion coupled multiangle light scattering (SEC-MALS). The data clearly demonstrated a single species in solution with an apparent molecular weight of 46 kDa, or a dimer of two EndoS1 monomers (23 kDa) (FigureS5A). The AlphaFold model^33^ provided in Uniprot for EndoS1 does not predict that the protein is a dimer in solution, however it does accurately predict the EndoS1 monomeric structure. When overlayed with each protein chain from the crystal structure individually, the AlphaFold model shows excellent agreement (Figure S5B). Furthermore, the two EndoS1 crystallographic monomers are identical (Figure S5B).

**Figure 3:**
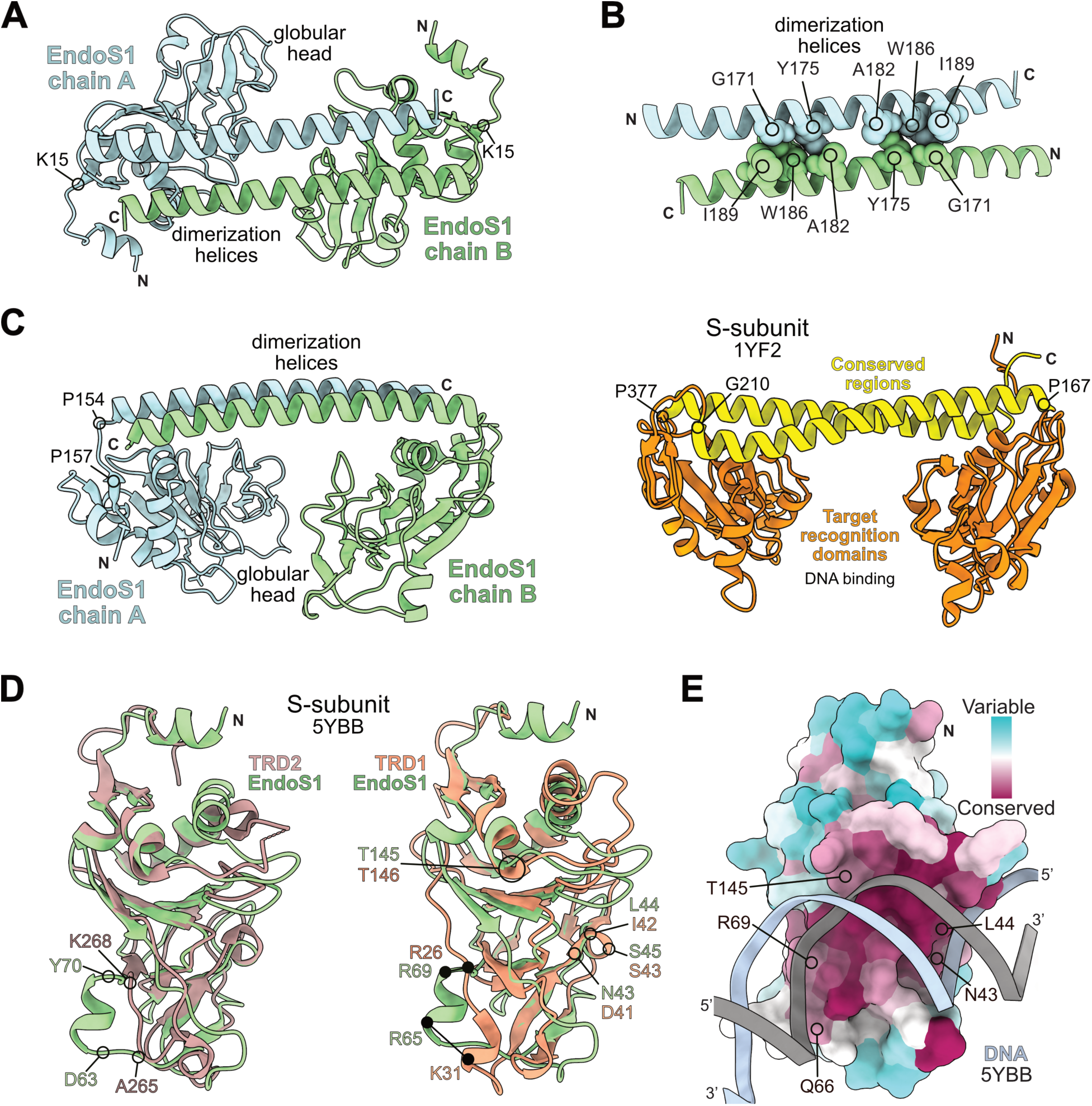
X-ray crystal structure of EndoS1. **(A)** Top-down view of EndoS1. The dimeric structure is labeled by dimerization helices and globular head with termini indicated. Residue K15 marks the start of the globular head domain fold. **(B)** Knob-in-hole packing of dimerization helices that likely stabilize the EndoS1 oligomer. **(C)** 90-degree rotation of EndoS1 from **(A)** compared to a DNA binding S-subunit (PDBid 1YF2). The EndoS1 dimerization helices are analogous to the S-subunit conserved regions and the EndoS1 globular heads are analogous to the S-subunit TRDs. The proline bounded loop linking the EndoS1 globular head and dimerization helices is indicated. The proline and glycine bounded loop linking TRD2 to the conserved region helix in S-subunit 1YF2 is also shown. **(D)** Structural overlay of the EndoS1 globular head (green) with both TRD2 (brown) and TRD1 (orange) from an S-subunit in complex with DNA (PDBid: 5YBB). EndoS1 contains an insertion helix residues D63 to Y70 relative to both TRDs. Left: The point where the EndoS1 helix is inserted compared to TRD2 is marked at TRD2 residues A265 and K268. Right: Essential residues in the S-subunit for DNA binding are indicated on TRD1. The residues in EndoS1 at the same positions as TRD1 DNA binding residues are also labeled. **(E)** Molecular surface of EndoS1 colored by residue conservation after structural alignment to a DNA-bound TRD from PDBid 5YBB. Residues critical for binding DNA are indicated. Residue conservation was determined by the server Consurf.

**Table 1:** Data collection and refinement statistics for EndoS1 and an EndoS1:Prx complex.

|  | EndoS1 | EndoS1:Prx complex |
| --- | --- | --- |
| **Data Collection** |  |  |
| Wavelength (Å) | 1.181 | 1.5418 |
| Space group | P121 | P61 |
| Cell dimensions |  |  |
| *a*, *b*, *c* (Å) | 82.36 43.87 85.84 | 67.49 67.49 232.21 |
| $\alpha$, $\beta$, $\gamma$ (°) | 90 118.67 90 | 90 90 120 |
| Resolution (Å) | 43.87-2.9 (3.19-2.9) | 33.75-2.69 (2.786-2.69) |
| Total reflections | 40593 (6573) | 1793759 (17965) |
| Unique reflections | 12014 (2947) | 16490 (1580) |
| CC(1/2) | 0.999 (0.940) | 0.998 (0.652) |
| *R*<sub>merge</sub> | 0.057 (0.464) | 0.141 (0.872) |
| *R*<sub>pim</sub> | 0.141 (0.872) | 0.045 (0.308) |
| *I*/σ*I* | 9.9 (2.0) | 13.3 (2.2) |
| Completeness (%) | 97.91 (97.32) | 99.47 (95.82) |
| Redundancy | 3.4 (3.4) | 10.8 (8.7) |
| **Refinement** |  |  |
| *R*<sub>work</sub> / *R*<sub>free</sub> (%) | 0.222/0.281 | 0.228/0.265 |
| Average B-factors (Å<sup>2</sup>) | 83.0 | 61.28 |
| Protein | 84.14 | 61.57 |
| Ligands | 112.34 | 58.15 |
| Water | 53.58 | 52.25 |
| No. atoms | 3141 | 4088 |
| Protein | 3086 | 3948 |
| Ligands | 5 | 27 |
| Water | 50 | 123 |
| Rms deviations |  |  |
| Bond lengths (Å) | 0.015 | 0.003 |
| Bond angles (°) | 2.1 | 0.53 |
| Ramachandran plot (%) |  |  |
| Total favored | 94.07 | 97.51 |
| Total allowed | 5.41 | 2.29 |
| PDB code | 8VSQ | 8VSR |
| Statistics for the highest-resolution shell are shown in parentheses |  |  |

As EndoS1 is annotated as a type I restriction endonuclease S-subunit (Figure 1), the crystal structure was compared to a known S-subunit from *Methanocaldococcus jannaschii* (PDBid 1YF2)^30^ (Figure 3C). When comparing EndoS1 to an S-subunit their overall structures are remarkably similar. The globular heads of EndoS1 are homologous to the DNA-binding target recognition domains (TRDs) of 1YF2 (orange), and the dimerization helices match closely to the conserved regions (CR) (yellow) of the true S-subunit (Figure 3C and Figure S5C). Additionally, the globular heads and TRDs in both structures are linked to the helical regions by proline bounded loops (EndoS1 P154-157, and 1YF2 P167, P377). This allows the S-subunit flexibility in the TRD relative to the CR helices when interacting with DNA and the M-subunit^31,32^. Although globally structurally similar to an S-subunit, EndoS1 has a critically important difference. S-subunits are one protein chain while EndoS1 is a homodimer. In fact, based on structural and domain map comparisons the EndoS1 monomer appears to be half of an S-subunit (Figure S5C). Furthermore, S-subunits contain two structurally different DNA binding domains that recognize non-palindromic sequences (Figure S5C-D)^30^. In contrast, since EndoS1 is a homodimer the protein has potentially two identical DNA binding domains. A dimer of identical DNA binding domains is characteristic of transcription factors^20^.

Further comparison between the globular head of EndoS1 and the S-subunit TRDs highlight additional similarities. As shown in Figure 3D and Figure S5C, the overall structure of the EndoS1 globular head is highly similar to both TRD1 and TRD2 of an S-subunit. Specifically, Figure 3D shows the EndoS1 globular head domain aligned to TRDs of an S-subunit in complex with DNA from *Caldanaerobacter sbuterraneus subsp. tengcongesis* (PDBid 5YBB)^32^. When comparing EndoS1 to TRD1, we observe that residues necessary for interaction with DNA in the S-subunit:DNA co-crystal structure are conserved exactly in EndoS1, except for the positions of R26 and K31 in 5YBB^32^. Additionally, when structurally aligned to an S-subunit bound to DNA, the bound DNA appears to slot within a binding surface on the EndoS1 globular head without significant clashes (Figure 3E). Moreover, when residue conservation is plotted on the molecular surface of EndoS1 the aligned DNA interaction surface is highly conserved. Taken together, the structural data shows that EndoS1 is a DNA binding protein and that the globular head contains a DNA binding cleft.

Despite the high similarity of the EndoS1 globular head to S-subunit TRDs, there is a critical structural difference. Relative to TRDs, the globular head of EndoS1 contains an apparent ‘insertion-helix.’ As shown in Figure 3E and Figure S5C, EndoS1 residues D63 to Y70 form a small ɑ-helix that juts out from the core TRD-like fold. For example, relative to 5YBB TRD2 the same region is an extended loop from residues A265 to K268 which in part serves to connect other secondary structure elements. Interestingly, the EndoS1 insertion-helix contains two residues (R65 and R69) that likely participate in DNA binding. These EndoS1 residues are close in position structurally to TRD1 DNA binding residues R26 and K31, despite being distant in the sequence alignment (Figure 3D-E and Figure S5D). Given the positional divergence of these DNA binding residues in EndoS1 relative to an S-subunit, we hypothesize that the EndoS1 insertion-helix is important for recognition of a specific DNA sequence. Or at least DNA sequences different than those bound by canonical S-subunits.

### Prx binds directly to the EndoS1 insertion-helix

To understand how Prx binds EndoS1, we pursued a co-crystal structure of an EndoS1:Prx complex. We were able to obtain diffracting crystals of Prx bound to an N-terminal truncated version of EndoS1 that lacked the first 9 amino acids (Figure 4 and Table 1). As shown in Figure 4A, Prx binds EndoS1 with a stoichiometry of 1 to 1 with one Prx bound to each EndoS1 chain in the crystal structure. Prx binds exclusively to the globular head of EndoS1 at a surface that faces outwards relative to the dimerization helices. Examining the EndoS1:Prx interaction surface in detail, we observe that Prx makes extensive interactions with the EndoS1 insertion-helix (residues D63 to Y70) (Figure 4B). This includes residues R65 and R69 which are likely required for interaction with DNA (Figure 3C-D, and Figure 4B). Furthermore, the EndoS1:Prx binding surface is primarily electrostatic in nature and stabilized by an extensive network of hydrogen bonds and salt-bridges. The EndoS1 binding surface has a net positive charge indicative of many DNA-binding proteins and Prx has a complementary electronegative surface (Figure S6A). Additionally, the observed binding mode with EndoS1 is extremely similar to how Prx binds the DBD of ComR^22^. In fact, many of the same residues that Prx uses to bind ComR also participate in binding to EndoS1. These include Prx residues E6 and D32 that when substituted to alanine (E6A and D32A), disrupt the ability of Prx to bind ComR (Figure 4B)^22^.

**Figure 4:**
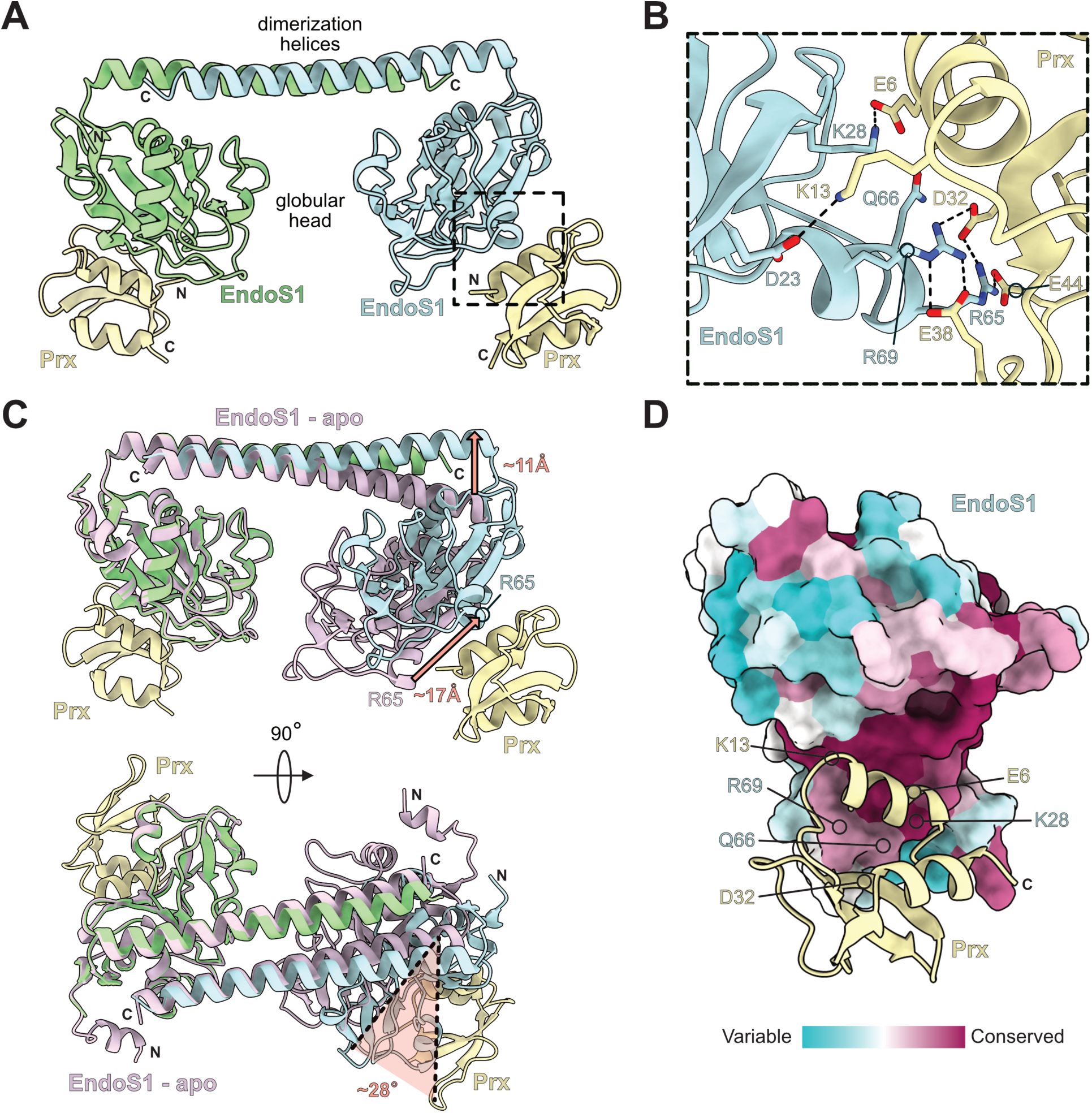
X-ray crystal structure of Prx bound to EndoS1. **(A)** Overall EndoS1:Prx complex structure showing Prx (pale goldenrod) binds each EndoS1 monomer (green, light blue) at the globular head. EndoS1 dimerization helices and globular heads are labeled. The EndoS1:Prx interaction surface is shown enclosed by dotted lines and is primarily composed of the EndoS1 insertion helix. **(B)** Detailed view of the EndoS1:Prx interaction. Hydrogen-bond and salt-bridge contacts are shown by dashed lines. **(C)** Structural alignment of EndoS1 (light purple) and the EndoS1:Prx complex structures. Conformational changes are highlighted by peach arrows and distances in Angstroms (top), and rotation relative to unbound EndoS1 (bottom). Residue R65 is shown to highlight the movement of the globular head and insertion helix. The rotation angle was calculated by the movement of R65 in unbound EndoS1 compared to EndoS1:Prx with Y117 as the rotation point. **(D)** Molecular surface of the EndoS1:Prx complex showing Prx binds in the conserved EndoS1 DNA binding cleft. EndoS1 is drawn as a molecular surface and colored by residue conservation as determined by the server Consurf. Prx is drawn in ribbon cartoon. EndoS1 residues (blue) and Prx residues (pale goldenrod) important for binding as shown in **(B)** are indicated on the structures.

The binding of Prx to EndoS1 also induces a significant conformational change in EndoS1. Compared to EndoS1 alone, binding of Prx causes the dimerization helices to flex outward by approximately 11Å (Figure 4C). This is also accompanied by a ∼28° rotation of a globular head out from under the dimerization helices, which results in a ∼17Å displacement of the Prx bound EndoS1 insertion-helix. Overall, the binding of Prx seems to force EndoS1 to shift from a rounded closed conformation to a more open conformation with the globular heads moved apart (compare the curvature of the dimerization helices, Figure 3A and Figure 4A, 4C). This conformation change seems to be limited to the dimerization helices and the globular head linker-regions as the globular heads do not change structure (Figure S6B). Moreover, the structure of Prx bound to EndoS1 is the same as both the Prx and ComR-bound crystal structures (Figure S6C)^21,22^.

Finally, binding of Prx to the EndoS1 insertion-helix allows Prx to slot directly into the EndoS1 DNA-binding cleft. Figure 4D shows the EndoS1:Prx complex with the molecular surface of the globular head colored by residue conservation. As clearly demonstrated, Prx is bound directly over several conserved EndoS1 DNA binding residues in the lower portion of the cleft. Furthermore, when DNA from the 5YBB crystal structure is overlayed onto EndoS1 as in Figure 3E, we observe significant structural clashes with bound Prx (Figure S6D). This shows that the interaction of EndoS1 with DNA would be incompatible with binding to Prx. Given the EndoS1:Prx co-crystal structure, functional analogy to Prx with ComR, and comparison to S-subunits, we conclude that Prx binds the EndoS1 insertion-helix to block the ability of EndoS1 to recognize DNA.

### Prx makes identical residue and structural contacts to EndoS1 and ComR

The EndoS1:Prx complex shows that Prx uses many of the same residues to bind EndoS1 as it does to bind ComR (Figure 4B). As such, we also tested the binding of EndoS1 to several Prx residue variants created from our previous studies. These variants included PrxE6A, PrxE9A, and PrxD32A, which we show disrupt binding to ComR (Figure 5)^22^, and PrxF31W which we have used as a spectroscopic probe to monitor Prx folding (Figure S7)^34^. Similar to ComR, PrxE6A and PrxD32A disrupt complex formation as assayed by SEC while PrxF31W still binds EndoS1 (Figure 5A and Figure S7). Interestingly, PrxE9A remains able to bind EndoS1 although PrxE9A does not interact with ComR^22^. Overall, these data both validate the complex observed in the EndoS1:Prx co-crystal structure (Figure 4), and demonstrate that Prx primarily uses the same molecular surface to contact EndoS1 as it uses to bind ComR.

**Figure 5:**
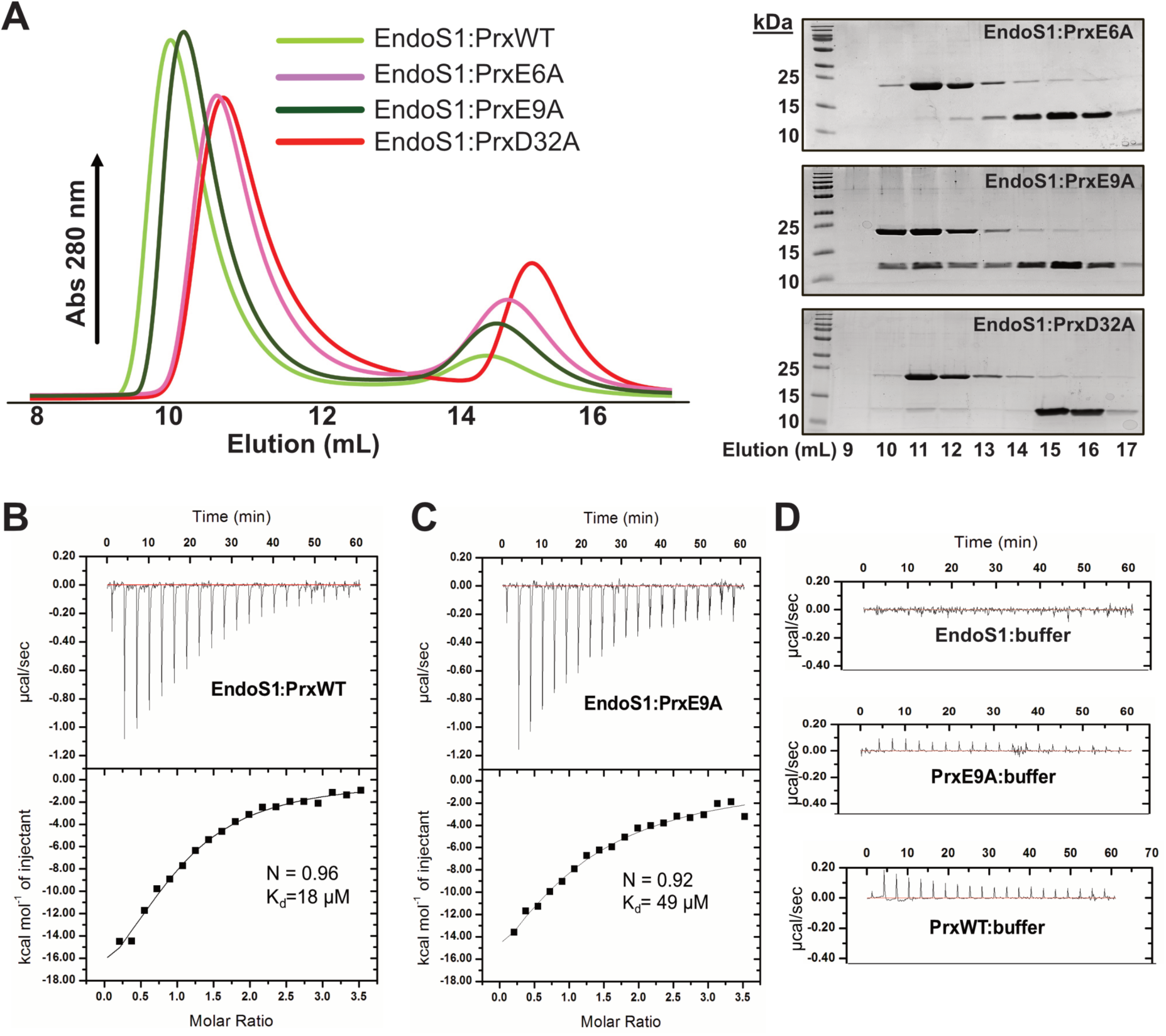
Prx variant binding assays with purified EndoS1. **(A)** SEC binding assay of EndoS1 with different Prx variants. EndoS1 was mixed with Prx variants separately to assay complex formation by SEC. The expected shift in elution volume for complex formation with wild type Prx (green) was absent with PrxE6A (light purple) and PrxD32A (red). PrxE9A was still able to form a complex with EndoS1. The adjacent panels show SDS-PAGE gels of the representative volumes from each SEC run. SEC fractions collected from the respective elution volumes in mL are shown. **(B)** Isothermal titration calorimetry (ITC) experiment of EndoS1 with Prx. Prx binds EndoS1 with an affinity of 18 µM and a stoichiometry of 1:1. **(C)** ITC experiment of EndoS1 with PrxE9A. PrxE9A binds EndoS1 with a reduced affinity of 49 µM and a stoichiometry of 1:1. **(D)** ITC controls of buffer titrated into EndoS1, and both PrxWT and PrxE9A titrated into buffer.

To further probe the interaction of EndoS1 with Prx, we also assayed their binding by isothermal titration calorimetry (ITC). As shown in Figure 5B, Prx binds EndoS1 with a stoichiometry of 1:1 as expected and a dissociation constant (K_d_) of 18 +/- 3 μM. The calculated ΔH and ΔS values were -24.8 +/- 2.2 kcal/mol and -61 cal/mol/deg, respectively. In contrast, ITC experiments of Prx with ComR revealed a K_d_ of 390 nM, which is essentially an order of magnitude tighter binding^22^. This demonstrates that the interaction of Prx with EndoS1 is significantly weaker than that with ComR. Additionally, as PrxE9A binds EndoS1 but does not bind ComR, we also tested this variant by ITC. As shown in Figure 5C, the apparent K_d_ is 49 +/- 11 μM, with calculated ΔH and ΔS values of -36.8 +/- 11.3 kcal/mol and -104 cal/mol/deg, respectively. The higher K_d_ may suggest a possible weakening of the EndoS1:Prx complex by PrxE9A. However, given typical protein preparation variability a less than 3-fold change in observed K_d_ could fall within experimental error. Note that ITC experimental controls for Figure 5B-C are shown in Figure 5D. Although Prx makes many identical contacts with ComR and EndoS1, there are differences such as PrxE9 that may help explain the large variation in binding affinity. Prx binds both ComR and EndoS1 using approximately the same molecular surface despite these proteins adopting completely different protein folds. Specifically, ComR is an HTH-fold DBD linked to a tetratricopeptide repeat (TPR) domain^19,20^ and EndoS1 is a dimer of S-subunit-like globular head TRDs (Figure 3). As such, we attempted to examine ComR and EndoS1 structurally for any common molecular elements. Given that Prx adopts the same conformation bound to ComR as it does EndoS1 (Figure S6C) we aligned a ComR:Prx complex (PDBid 7N10) to the EndoS1:Prx complex. As shown in Figure 6A, the results are striking. When overlayed, the EndoS1 insertion-helix and the DNA-binding helix of the ComR-DBD match almost exactly. This includes the conserved ComR DNA binding residues R33, Q34, and R37 which are also present in the EndoS1 insertion-helix at positions R65, Q66, and R69. Furthermore, the sidechains of both R37 (ComR) and R69 (EndoS1) adopt exactly the same conformation so as to contact residue D32 in Prx. Again, substitution of PrxD32A inhibits binding to both EndoS1 (Figure 5) and ComR^22^. To further verify the importance of the EndoS1 insertion-helix, both EndoS1 Q66 and EndoS1 R69 were substituted with alanine. Notably, EndoS1R69A failed to bind Prx confirming the importance of the arginine-aspartate H-bond in the stability of the EndoS1:Prx complex (Figure 6B). Taken together, Prx not only uses the same molecular surface to bind both ComR and EndoS1 but recognizes a conserved positively charged DNA binding helix present in both proteins.

**Figure 6:**
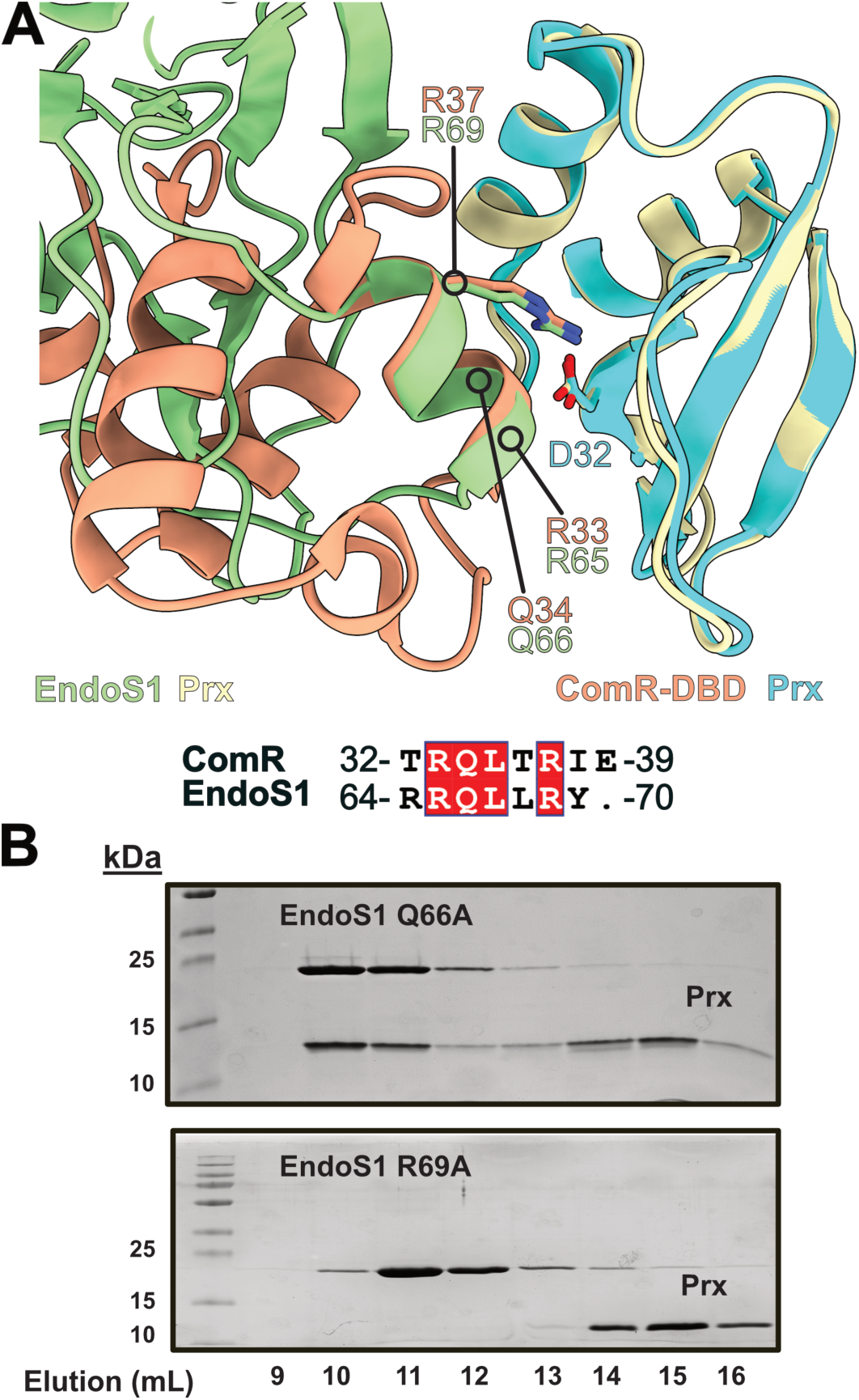
Structural comparison of EndoS1:Prx and ComR:Prx. **(A)** Structural alignment of EndoS1:Prx (green:pale goldenrod) with ComR:Prx (PDBid 7N10) (orange:cyan). The ComR structure consists only of the DNA binding domain (DBD). Equivalent residues between the EndoS1 insertion helix and the ComR DNA-binding residues are highlighted on the structure. A sequence alignment of the EndoS1 insertion helix with the ComR-DBD is shown and colored by residue identity (red box). Prx residue D32 that is critical for binding both EndoS1 and ComR is also shown. **(B)** EndoS1 variant SEC binding assay with Prx. Coomassie stained SDS-PAGE gels are shown of EndoS1Q66A and R69A with wild-type Prx. For all SDS-PAGE gels, SEC fractions collected from the respective elution volumes in mL are shown. SEC chromatograms are shown in Figure S8

### The binding mechanism of Prx to JM3 is different from Prx with EndoS1 and ComR

As we have explored the molecular details of the interaction of Prx with both EndoS1 and ComR, we employed a similar analysis with the phage protein JM3. As Prx and JM3 form a stable complex by SEC (Figure 2) we also attempted to obtain crystals of the complex. To date, diffracting crystals have not been obtained with the current materials. However, as AlphaFold2 predicts JM3 to adopt an HTH-DBD (Figure S9A) we aligned the predicted JM3 structure to a ComR:Prx complex (PDBid 7N10) (Figure 7A). JM3 and the ComR-DBD align extremely well indicating that both proteins adopt the same fold of an HTH-DBD. However, when comparing the ComR DNA binding helix with the predicted JM3 model we observe no conservation. Additionally, a sequence alignment of the ComR DBD helix, the EndoS1 insertion helix, and the equivalent secondary structure in JM3 shows very little homology (Figure 7B). Importantly, the critical Prx binding residue R37/R69 (ComR/EndoS1) is an alanine in JM3 (A57). Moreover, the molecular surface of the JM3 predicted model contains only two basic residues. This is in stark contrast to EndoS1 and ComR, which both have very electropositive surfaces (Figure S6A)^22^. Given these observations, the current structural data cannot explain how Prx binds to JM3.

**Figure 7:**
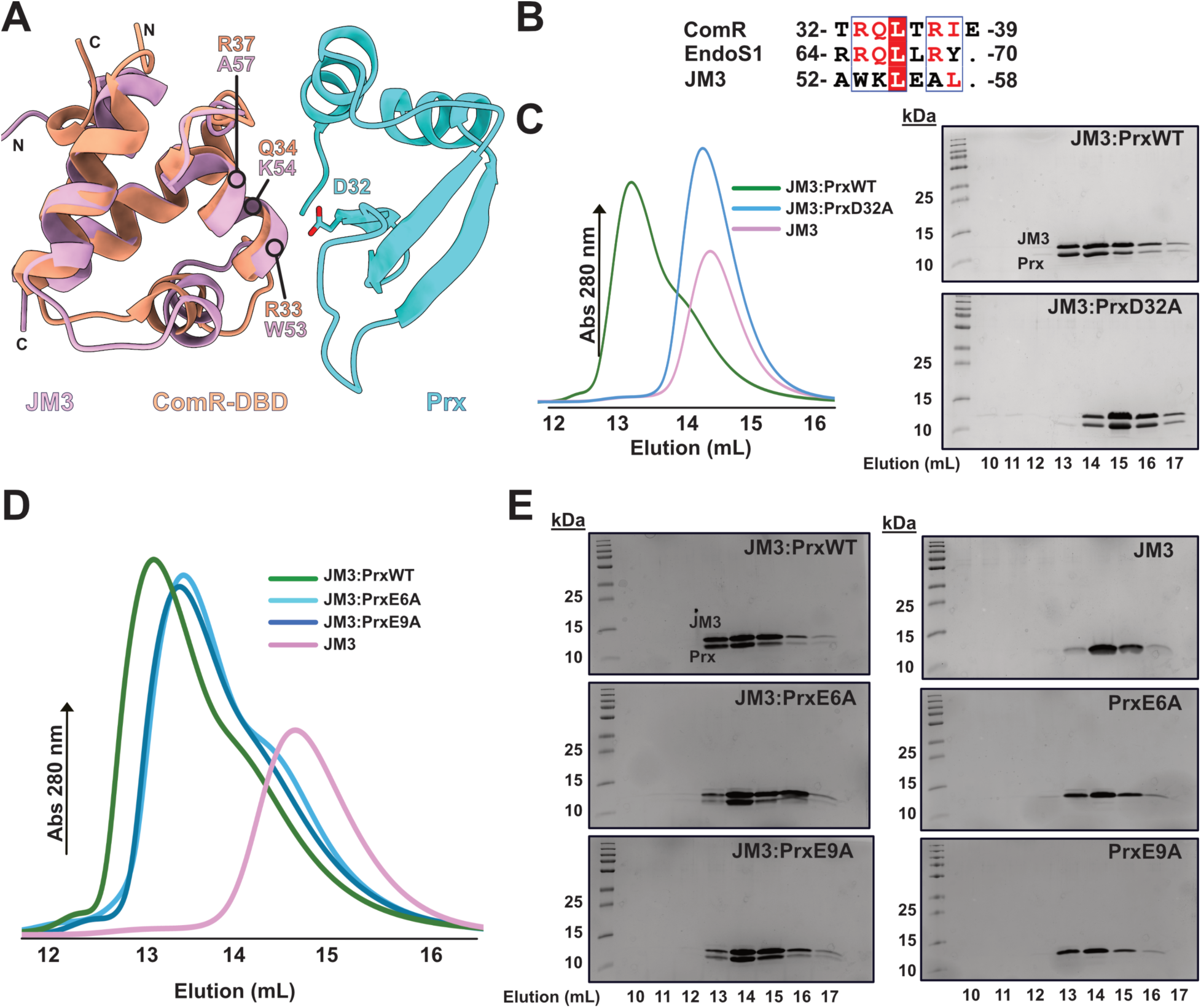
Molecular interaction of Prx with JM3. **(A)** Overlay of a JM3 AlphaFold model (violet) with a ComR:Prx cocrystal structure (PDBid 7N10) (orange:cyan). ComR residues important for binding Prx are shown with equivalent position residues in JM3. Prx D32 is also shown. **(B)** Sequence alignment of the ComR DNA binding residues, the EndoS1 insertion helix, and the structurally aligned helix of JM3. **(C)** SEC binding assay of JM3 with PrxD32A. PrxD32A disrupts binding with JM3 which can be observed from the absence of shift in elution peak (pink) compared to JM3:PrxWT (green). The SEC trace of JM3 alone is shown in blue. The adjacent panels show Coomassie stained SDS-PAGE gels of the representative volumes from the JM3:PrxWT and JM3:PrxD32A SEC runs. SEC fractions collected from the respective elution volumes in mL are shown. **(D)** SEC binding assays of JM3 with Prx variants E6A (light blue) and E9A (dark blue). **(E)** Coomassie stained SDS-PAGE gels of complexes (left) and proteins alone (right).

To understand how Prx binds JM3, we also performed binding assays of JM3 with Prx variants. As shown in Figure 7C, PrxD32A does not bind JM3 as assayed by SEC. Namely, a shift of the proteins to a larger molecular size in solution is not observed. We also tested the binding of JM3 to the Prx variants PrxE6A and PrxE9A (Figure 7D-E). Both variants appear to bind Prx, although the solution properties of the variant complexes appear differ from the wild-type JM3:Prx complex. Specifically, the elution volumes of the variant complexes appear to have shifted to a smaller apparent size as demonstrated by a later elution volume (Figure 7D). In addition to the critical D32 residue, the data shows that Prx uses a different set of residues to contact ComR (E6, E9, and D32), EndoS1 (E9 and D32), and JM3 (D32).

Although Prx requires D32 to bind JM3 in our assay conditions, JM3 lacks the conserved arginine that contacts PrxD32 found in both ComR and EndoS1. As such, it is unknown how PrxD32 contributes to binding JM3. To address this, we attempted to model a JM3:Prx complex using AlphaFold2. The resulting prediction was of high confidence (Figure S9B) and predicted an overlapping but different contact surface from that shown in Figure 7A. AlphaFold2 predicts that Prx still makes contacts with the ComR aligned helix (Figure 7A), but Prx is shifted on the JM3 surface (Figure S9C). The prediction pushes PrxD32 by about ∼5.9Å away from the ComR aligned helix in JM3 (Figure S9C). In this new position, PrxD32 could potentially make H-bond or salt bridge contacts with other residues or the JM3 mainchain (Figure S9C). To determine the molecular contacts, an experimental structure will be required. Although the AlphaFold prediction of the JM3:Prx complex is of high confidence, Prx is an intrinsically disordered protein^34^. Specifically, our previous work has shown that Prx is in equilibrium with a disordered state and a globular fold. Moreover, this unbound Prx globular fold is different from the structure Prx finally adopts when bound to either EndoS1 or ComR^34^. As such, the conformation of Prx bound to JM3 could be different from the known crystal structures and therefore unpredictable by AlphaFold. Although Prx utilizes the same residues to contact JM3 as it does EndoS1 and ComR, the details of the molecular interaction with JM3 are significantly different.

### Several Prx binding partners contain a conserved helical motif

The biochemical data shows that Prx interacts with at least two of its binding partners by the same mechanism. Specifically, Prx contacts ComR and EndoS1 by binding a highly conserved ɑ-helix present in both proteins (Figure 6). Given this observation, we asked if the same positively charged helix was also found in other putative Prx binding partners. Using the experimental ComR:Prx and EndoS1:Prx structures as a guide, we used AlphaFold2^33^ to model Prx in complex with select proteins from the mass-spectrometry pull-down data (Figure 1, Figures S1-3). Given the extensive list, we focused on the putative binding partners selected to be shown in Figure 1, those hits predicted to be DNA binding proteins, and small proteins amenable to modeling.

We first started with SpyM3_1346 because it contains the same HTH-fold subtype as ComR and is a putative lambda repressor (Figure 1). Lambda repressors are transcriptional regulators that control the transition between the phage lysogenic and lytic cycles^35^. These proteins contain HTH-DBD domains that oligomerize to bind DNA and can also bind RNA polymerase^36–38^. Alphafold2 produced an extremely confident model of a SpyM3_1346:Prx complex using SpyM3_1346 residues 1-58 (Figure S10A, Figure 8A). When compared to a ComR:Prx structure (PDBid 7N10), the predicted SpyM3_1346:Prx complex aligns very closely. The conformation of Prx is identical and SpyM3_1346 adopts an HTH-DBD fold similar to the ComR-DBD. Additionally, the ComR DNA binding helix matches almost exactly to a helix in SpyM3_1346 that is predicted to contact Prx. Moreover, the SpyM3_1346 helix includes residues R27 and R31 which match R33 and R37 in ComR, respectively. Finally, R31 is predicted to bind PrxD32 similar to R37 in ComR and thus R69 in EndoS1. Given the putative phage regulation function of SpyM3_1346 and its molecular conservation with ComR, the predicted AlphaFold model is likely an accurate representation of the real Prx bound complex. In line with this, Prx likely binds SpyM3_1346 to also block its interaction with DNA.

**Figure 8:**
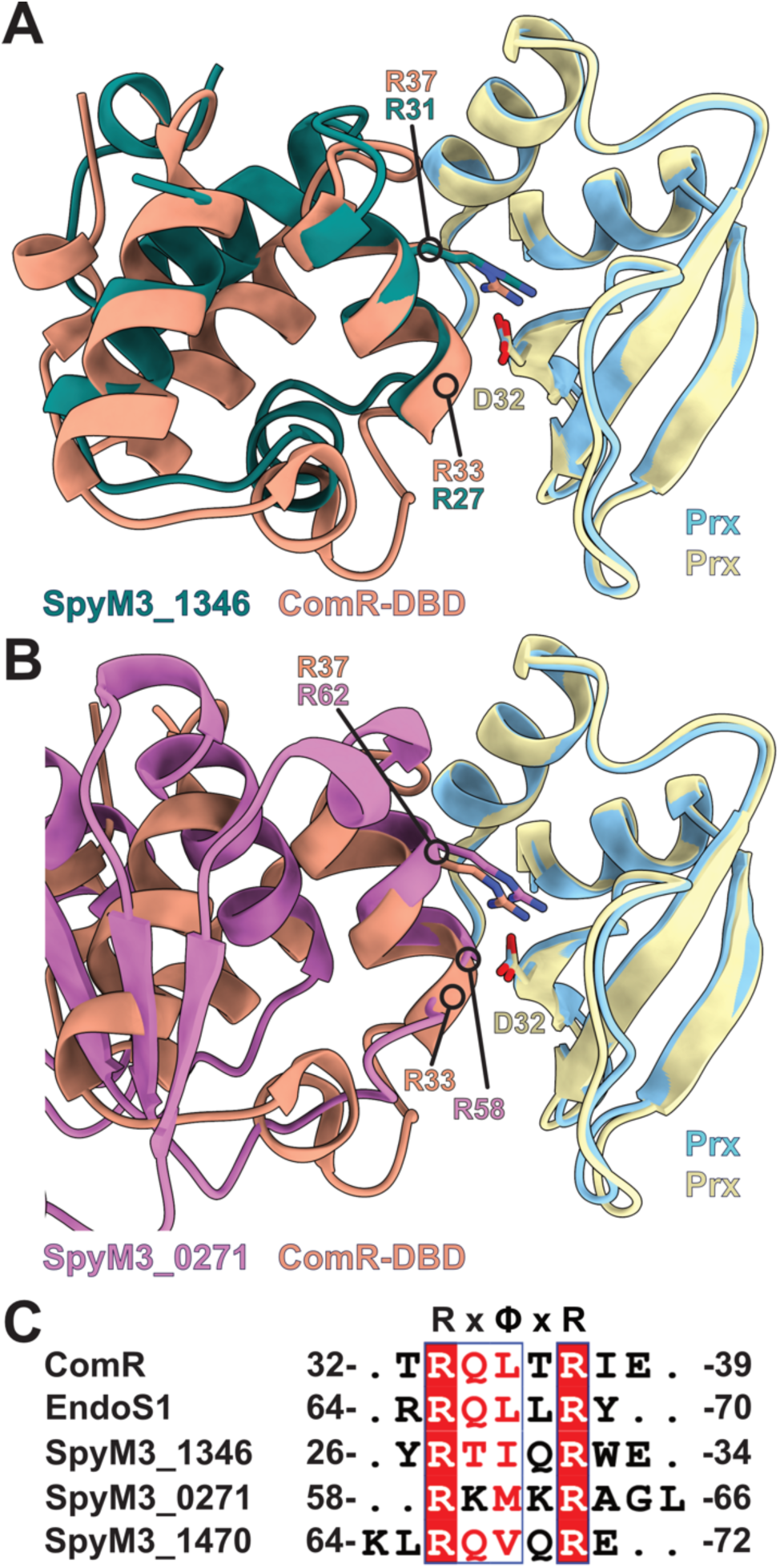
Structural models of Prx protein complexes reveals a conserved binding motif. **(A)** ComR:Prx (PDBid 7N10) (orange:pale-goldenrod) structural aligned to a SpyM3_1346:Prx complex predicted by AlphaFold (teal:cyan). The critical interaction between ComR R37 and Prx D32 is preserved in SpyM3_1346 at residue R37 and Prx D32. SpyM3_1346 is a helix-turn-helix fold highly similar to ComR **(B)** ComR:Prx (PDBid 7N10) (orange:pale-goldenrod) structure aligned to a SpyM3_0271:Prx complex predicted by AlphaFold (violet:cyan). The critical interaction between ComR R37 and Prx D32 is preserved in SpyM3_0271 at residue R62 and Prx D32. SpyM3_0271 does not contain a HTH domain and is structurally different from both ComR and EndoS1. **(C)** Sequence alignment of the Prx interaction surfaces from experimentally determined complexes (ComR, EndoS1) and AlphaFold predicted complexes (SpyM3_1346 and SpyM3_0271). SpyM3_1470 contains the Prx interaction helix (Figure S10) but AlphaFold was unable to predict the complex. Residues are colored by sequence identity (red box) and homology (red letter) by the server Espript3 (https://espript.ibcp.fr/ESPript/ESPript/). The consensus motif is listed above the sequence, where x is any residue and <Ι is hydrophobic.

Another successful AlphaFold prediction was the putative tRNA-methyltransferase SpyM3_0271 (Figure 1). Full-length SpyM3_0271 modelled with Prx produced a moderate-to-high-confidence complex (Figure S10A). Additionally, as shown in Figure 8B SpyM3_0271 contains the same helical motif as ComR, EndoS1, and SpyM3_1346. When the predicted SpyM3_0271:Prx complex is aligned to a ComR:Prx complex, we observe that SpyM3_0271 contains arginine residues at the same critical positions for interaction with Prx. This includes residue R62 which adopts the same conformation as ComR residue R37 for binding PrxD32 (Figure 8B). SpyM3_0271 is annotated as a 2’-O-methyltransferase that modifies cytidine at the wobble position in tRNA. The reaction is specifically carried out using S-adenosyl-L-methionine as a substrate and is thought to promote RNA stability and folding^39^. The Prx binding helix in SpyM3_0271 is distal from the enzymatic active site. However, this helix has been shown to change conformation upon binding its substrate in closely related structures^40,41^. Additionally, the putative Prx binding surface does not appear to clash with the tRNA methyltransferase dimerization site. From the available 2’-O-methyltransferase experimental structures and the SpyM3_0271 model, it is currently unclear if Prx affects tRNA binding. As such it is unknown how Prx may affect the biochemical activity of SpyM3_0271.

SpyM3_1470 is a small protein of unknown function that was pulled-down by Prx from both XIP and mitomycin C induced lysates (Figure S1-2). The protein is a helical protein with a DUF896 annotation. DUF896 proteins such as YnzC are thought to oligomerize for DNA binding, but their exact biological function is unknown^42^. Although AlphaFold did not confidently predict a SpyM3_1470:Prx complex, the protein was small enough to manually observe the presence of a putative Prx binding helix (Figure S10B). Additionally, many DUF896 proteins are part of the bacterial SOS response to DNA damage and are under the control of the transcriptional repressor LexA. Notably, LexA represses the bacterial SOS response and is directly involved in the lysogenic/lytic life-cycle switch of many phages^43,44^. The switch to the lytic cycle for the prophage is induced when LexA and a phage repressor are cleaved by RecA^44,45^. As Prx pull-downs the putative phage repressor SpyM3_1346 (which contains the Prx binding helix), an SOS response protein (SpyM3_1470), and RecA (SpyM3_1800, Figure 1) we conclude that Prx plays a direct role in the lysogeny/lytic life-cycle decision of GAS. However, the mechanism and exact role of Prx in this metabolic process remains to be determined.

Finally, given our experimental data (Figures 3-5) and AlphaFold predictions (Figure 8, FigureS10) we asked if the Prx binding helix has a defined sequence motif. A multisequence alignment of all discovered Prx binding helices is shown in Figure 8C. As clearly demonstrated, there is in fact a conserved sequence motif. The Prx binding consensus sequence is RxΦxR, where Φ is any aliphatic hydrophobic residue. Additionally, the motif is net positive with polar residues at each x position with Q highly represented. However, it must be noted that Prx still binds JM3 which does not have this consensus sequence (Figure 7). Furthermore, the putative Prx binding partner SpyM3_0977 contains an HTH-domain of nearly identical in fold to both JM3 and ComR, but also lacks the Prx binding helix (Figure 1). Similar to JM3, AlphaFold predicts a shifted Prx binding mechanism relative to ComR:Prx (Figure S9 and Figure S10C-D). Given these observations, the presence of the Prx binding helix (RxΦxR) is highly predictive of Prx binding but the absence of the helix does not preclude interaction. Subsequently, this implies that Prx may adopt different conformations, or at least can recognize varied molecular surfaces, dependent upon its interaction partner.

## DISCUSSION

In this study we have shown that Prx interacts with numerous GAS host and bacteriophage proteins, with the general theme of interacting with DNA-binding proteins to block their interaction with DNA. Specifically in this study we have shown that Prx binds directly to the restriction endonuclease S-subunit-like GAS protein EndoS1 (Figure 2). Moreover, Prx recognizes an ‘insertion-helix’ found in the DNA binding surface of EndoS1 that is not found in canonical S-subunits (Figure 3-4). The effect of Prx binding is clearly to block the ability of EndoS1 to bind DNA. We also found that although the protein fold of EndoS1 and ComR are completely unrelated, the EndoS1 ‘insertion helix’ is found exactly in the DBD of ComR (Figure 6). Furthermore, the shared Prx binding helix and its sequence motif (RxΦxR) are found in several other putative Prx binding partners (Figure 8). Additionally, we have shown that Prx directly binds the phage protein JM3 (Figure 2). Although JM3 has the same fold as the DBD of ComR, it lacks the Prx binding helix (Figure 7-8). This shows that Prx has the ability to interact with various proteins with different molecular surfaces. Overall, Prx is a multifunctional protein that binds to various transcriptional regulators and other DNA-binding proteins. Given this, Prx appears to be a metabolic regulator of its bacterial host with direct influence on DNA processing and the lysogeny/lysis decision of the phage.

In addition to being pulled-down from all cell lysates, EndoS1 was also of interest due to its annotated biological function of being an S-subunit (Figure 1). This immediately raised the hypothesis that EndoS1 may be part of a type I restriction endonuclease complex that is specific for *prx* harboring phages. However, as shown by the EndoS1 crystal structure it varies significantly from a canonical S-subunit which casts some doubt on this hypothesis (Figure 3). Namely, EndoS1 is a dimer of equivalent TRDs for DNA binding whereas S-subunits are one polypeptide with two different TRDs (Figure 3 and Figure S5)^31,32^. Furthermore, GAS has a canonical S-subunit encoded together with a methylation subunit. These proteins are HsdS (SpyM3_1643) and HsdM (SpyM3_1644), respectively. In our hands we were not able to show conclusively that EndoS1 binds HsdM by SEC (Figure S11A-B). There is minor evidence of a change in elution volume when EndoS1, Prx, and HsdM are added together but the data are not strong enough to claim binding. This is in contrast to HsdS and HsdM which readily for a stable complex (Figure S11C). For completeness, we also show that Prx does not bind the HsdS:HsdM complex (Figure S11D). We do note that adding EndoS1 to HsdM did cause extensive precipitation, but that behavior alone is not enough to prove a biologically relevant interaction. Given that EndoS1 may not bind HsdM and that the EndoS1 structure diverges significantly from known S-subunits, EndoS1 is likely not an S-subunit. This is supported by that EndoS1 is encoded distal from type-I restriction genes (Figure S4), and that EndoS1 is a dimer of identical DNA binding domains (Figure 3). The structural configuration of EndoS1 is instead characteristic of a transcription factor or repressor^18,45,46^. Regardless, additional experimentation is required to discover the biological function and DNA recognition sequence for Endos1.

Another reason why EndoS1 was of interest is that its protein fold is completely unrelated to that of the previously discovered Prx binding partner ComR^19,21,22^. However, upon solving an EndoS1:Prx crystal structure we observed that EndoS1 has similar molecular properties to ComR. This includes a positively charged DNA and Prx interaction surface, in addition to an ɑ-helix (EndoS1 insertion-helix) that contacts Prx residue D32 (Figure 4 and Figure S6). We also observed that Prx induces significant conformational changes in EndoS1 (Figure 4), which is similar in effect to Prx on ComR^22^. Specifically, Prx binds the DNA-interaction surfaces of both EndoS1 and ComR while manipulating their conformations with the net-effect of inhibiting the ability of both EndoS1 and ComR to bind DNA.

Based on the observation that Prx binds to a charged helical structure in both ComR and EndoS1 (Figure 6), we searched for this structural motif in other putative Prx binding partners (Figure 1). As shown in Figure 8C, a conserved ɑ-helix binding motif can be found in at least 5 Prx binding partners. The consensus sequence contains 2 arginine residues spaced to be on the same face of an ɑ-helix to presumably interact with DNA. In between the arginine residues is an aliphatic residue on the opposite helix face to pack within a hydrophobic core (RxΦxR). Taken together, our structural and predictive modeling data strongly suggests that this Prx structural binding motif maybe present in many nucleic acid binding proteins. However, given that RxΦxR is a small motif a sequence-based search alone is likely to produce many false-positives. As such, any large scale-data mining should include a structural component. Although beyond the scope of this study, such genome-wide searches are of interest not only in GAS but in other Streptococcal species which harbor Prx. For example, Prx is also found in Group B Streptococcus.

Only 4 of the 5 proteins found to contain the Prx binding helix are DNA-binding proteins, with one of the hits annotated as a tRNA methyltransferase (SpyM3_0271) (Figure 1, Figure 8C). The GAS tRNA methyltransferase clearly contains the Prx binding helix but the biological reason for this interaction is unclear. Regardless, SpyM3_0271 was one of the most probable hits from the mass-spectrometry data. Moreover, Prx co-preciptated with several other methyltransferases suggesting a true interaction with biochemical relevance (Figure 1, Figures S1-3). Given that ribosomal and RNA binding proteins are common false-positives in bead-based co-precipitation experiments^26^, the significance of the interaction of Prx with tRNA methyltransferases requires additional validation.

In addition to binding bacterial proteins such as ComR and EndoS1, we have now shown that Prx also directly binds another phage protein. Prx forms a stable complex with the phage protein JM3, which is also an HTH DBD fold similar to ComR (Figure 2 and Figure 7). Our biochemical results show that Prx binds all three of its experimentally verified proteins using the conserved residue D32 (Figure 5 and Figure 7)^22^. However, JM3 lacks the Prx binding helix (Figure 7 and Figure 8). AlphaFold instead predicts that Prx recognizes a surface on JM3 different from the surface Prx binds on the ComR-DBD (Figure S9C-D). Likewise, AlphaFold makes a similar prediction with the putative Prx binding partner SpyM_0977 which also has a ComR-like DBD but lacks the Prx binding helix (Figure S10D). Furthermore, Prx co-precipitates with other DBD containing proteins such as SpyM3_0843 and SpyM3_1036 (BirA). The DBD-fold found in these proteins is not the same protein family as ComR and JM3, and the fold also appears to also lack the Prx binding helix motif (RxΦxR). We have previously shown that Prx is a highly dynamic protein and is in equilibrium between an intrinsically disordered ensemble and a compact globular fold. Moreover, this unbound Prx fold is distinct from the fold Prx adopts when it binds ComR or EndoS1^34^. Considering the apparent binding promiscuity of Prx and its dynamic properties, it is tempting to speculate that Prx may adopt a different conformation when it complexes with proteins that lack the RxΦxR binding helix. However, an experimental complex of Prx bound to JM3 or another binding partner will be required to test this hypothesis.

Like EndoS1, the biological function of JM3 is currently unknown. In contrast to EndoS1, Prx only co-precipitated with JM3 from lysates made from cells in the lytic cycle (+mitomycin C, Figure 1)^9^. This suggests that JM3 may participate in some aspect of the lysogeny to lytic transition, phage assembly, and/or be responsible for regulating phage gene expression. These hypotheses are supported in part due to the location of the JM3 gene (*spyM3_1246*). Specifically, JM3 is in a phage gene cluster that appears to function in DNA processing (Figure S4). These genes include a single-stranded DNA binding protein, a DNA recombination protein, RusA, and a Siphovirus Gp157 protein. Gp157 proteins increase resistance to bacteriophages^47,48^ and RusA family members resolve intermediate structures made during DNA repair and recombination^49,50^. In fact, RusA (SpyM3_1248) was also found to co-precipitate with Prx only from lytic cycle lysates (Figure 1). Taken together, this strongly suggests that JM3 plays a role in the regulation of phage DNA processing. The AlphaFold model of JM3 is a single DBD with an unpredicted 22 amino-acid tail (Figure S9), raising the hypothesis that JM3 may dimerize and bind DNA like a transcriptional repressor. Regardless, the biochemical function of Prx is likely to inhibit any regulatory effect of JM3.

Our results show that Prx plays a large role in GAS metabolism. We have demonstrated that Prx binds bacteria and phage proteins alike, most of which are transcriptional regulators. Importantly our biochemical data suggests that Prx has a role in the phage lysogenic/lytic cycle, DNA recombination, and potentially phage assembly. These biological activities are in addition to our past work showing that Prx inhibits both natural competence and quorum sensing^21,22^. Furthermore, these findings may help explain the drastically increased electrocompetence phenotype we have observed in a *prx* deletion strain^21^. Without Prx blocking the interaction of numerous DNA processing proteins with DNA, exogenous DNA can be more readily taken up and stabilized. Additionally, as EndoS1 is in a gene cluster with transporters the electrocompetence phenotype may also be related to a change in GAS membrane permeability. Overall, our work clearly demonstrates that the bacteriophage utilizes Prx as a diverse and multifunctional binding protein to control the biology of its GAS host.

Finally, we had previously hypothesized that Prx inhibits ComR to protect the phage genome from recombination^21^. Expanding this with our current findings, we now hypothesize that the overall biological function of Prx is to protect the genetic integrity of the phage genome (including the genetically linked toxin gene). Namely, the phage uses Prx to manipulate the DNA metabolism of the GAS host to favor the phage. Subsequently we propose that without Prx, phage genes such as the toxin gene would be more easily lost and the evolutionary disadvantage of GAS being a pathogen to its only host will be removed.

## MATERIALS AND METHODS

### Molecular cloning of protein expression constructs

Recombinant protein expression constructs created in this study included EndoS1 (SpyM3_0890), JM3 (SpyM3_1246), HsdS (SpyM3_1643), and HsdM (SpyM3_1644). All genes were PCR amplified using genomic DNA from strain MGAS315. Wild-type Prx and Prx residue variants were created previously^21,22^. EndoS1 and HsdM were restriction cloned into pET22b using NdeI and XhoI sites to add a C-terminal 6His-tag using primer pairs P140/P141 and P158/P159 respectively. Truncated EndoS1 lacking the first 9 amino acids was created by Q5 mutagenesis (New England Biolabs, NEB) by amplifying the full-length construct using primer pairs P154/P155. JM3 was amplified from MGAS315 genome using primer pairs P202/P203 and was placed in pGEX6P-1 between the BamHI and NotI sites to create a GST-JM3 chimera. For HsdS expression, both HsdS and HsdM were amplified with primer pairs P206/P152 and P207/P183 respectively and placed into pETDUET-1. HsdS and HsdM were cloned into multiple cloning site 1 with NdeI and XhoI and cloning site 2 with EcoRI and NotI, respectively. All EndoS1 variants were generated by Q5 mutagenesis. EndoS1 variant Q66A was generated by amplifying the full-length construct with primer pair P211/P212. EndoS1variant R69A was generated by amplifying the full-length construct with primer pair P213/P214. Ligated vectors were transformed into competent *Escherichia coli* BL21(DE3)-Gold for protein expression. A list of primers used is supplied in Table S1.

### Protein expression and purification of Prx, EndoS1, HsdS, and HsdM

Prx, EndoS1, HsdS, and HsdM and their variants were all purified by the same metal-affinity protocol outlined here. Note, additional details for Prx are described previously^22^. Each expression construct in BL21(DE3)-Gold cells was pre-cultured overnight in 30 mL of Lysogeny Broth (LB) broth with 100 μg /mL ampicillin. The following day they were transferred to 3 litres of fresh LB broth at a 1 to 100 dilution and grown at 37°C until an OD_600nm_ of 0.6. Protein expression was then induced by adding 1 mM IPTG and cells were allowed to grow overnight at 20°C. The cells were collected by centrifugation at 4200 rpm for 30 minutes and resuspended in 100 mL lysis buffer (50 mM Tris-HCl pH7.5, 500 mM NaCl, 25 mM imidazole). Before lysis, resuspended cells were treated with 1 mM phenylmethylsulfonyl fluoride (PMSF), 10 mM MgCl_2_, and a small amount of DNase I. The cells were then lysed under high-pressure using an Emulsiflex-C3 (Avestin). The cell lysate was cleared by centrifugation at 17,000 rpm (30,000xg) at 4°C for 30 minutes to remove insoluble material. The lysate supernatant was then passed over a nickel-NTA affinity gravity flow column (GoldBio) pre-prepared in lysis buffer. After binding the protein, the column was washed with at least 20x column volumes of lysis buffer before being eluted with 3x column volumes of elution buffer (50 mM Tris pH 7.5 500 mM imidazole). All buffers where chilled before use.

After metal-affinity chromatography each protein was then further purified by size-exclusion chromatography (SEC) using an AKTApure (Cytiva) and a HiLoad 16/600 Supderdex75 column (Cytiva) or HiLoad 16/600 Supderdex200 column. Before injection all proteins were concentrated using an Amicon centrifugal filter with either a 3 kDa or 10 kDa cut-off depending on molecular weight. All SEC purifications were performed at 4°C. The gel filtration buffer for Prx was composed of 20 mM Tris at pH7.5, 100 NaCl, and 1 mM beta-mercaptoethanol (β-Me). The gel filtration buffer for EndoS1, HsdM and HsdS was composed of 50 mM HEPES at pH 7.5, 150 mM NaCl and 1 mM β-Me. Pure fractions as assayed by Coomassie-stained SDS-PAGE gel were collected and concentrated by an Amicon centrifugal filter and flash frozen in liquid nitrogen for future use.

### Protein expression and purification of JM3

GST-JM3 was expressed in BL21(DE3)-Gold cells, with cells harvested and lysed as described for both Prx and EndoS1. However, for GST-JM3 the lysis buffer consisted of 50mM Tris pH7.5, 200 mM NaCl_2_, and 2 mM β-Me. For purification, the GST-JM3 containing bacterial supernatant was first passed over a Q-sepharose (Cytiva) gravity column equilibrated in GST lysis buffer. This step removes cellular debris, some proteins, and DNA. Next the pre-cleared lysate was allowed to flow over a gluthione-sepharose-4B (Cytiva) gravity column. The column was washed with at least 20x column volumes of GST wash buffer (50 mM Tris pH 7.5, 500 mM NaCl, 2 mM β-Me) and GST-JM3 eluted in 3x column volumes of GST elution buffer (50 mM Tris pH 7.5, 500mM NaCl, 20 mM glutathione). Following elution the GST-tag was removed by over-night cleavage at 4°C using a 1:50 (enzyme:protein) dilution of HRV-3C protease. During GST-tag digestion, the sample was dialyzed into GST lysis buffer. After the digestion was complete, the dialyzed material was passed back over a gluthione-sepharose-4B gravity column to remove GST and uncleaved GST-JM3.

Following affinity purification JM3 was also further purified by SEC using the same protocol as Prx and EndoS1. For JM3 the SEC buffer used was (20mM Tris pH 7.5, 100mM Nacl, 1mM β-Me). Finally, Pure fractions as assayed by Coomassie-stained SDS-PAGE gel were collected and concentrated by an Amicon centrifugal filter and flash frozen in liquid nitrogen for future use.

### Creation of GAS cell-lysates for Prx pull-down experiments

*S. pyogenes* stain MGAS315 was streaked directly from a freezer stock on trypticase soy agar (TSA II) plates supplemented with 5% sheep’s blood (Blood Agar, ThermoFisher). The cells were then allowed to grow for a minimum of 24 hours at 37 °C. Blood agar was used to detect GAS by beta-hemolysis and to verify that no contaminants were present. Colonies were picked and inoculated in 7.5 mL of pre-warmed (37 °C) chemically defined media (CDM)^16,51^ in a sealed screw-capped tube and allowed to grow overnight at 37 °C. Overnight cultures were diluted into fresh CDM at a 1:20 ratio in triplicate sets, and grown to an OD_600nm_ of 0.2. At this point media was supplemented with 50 nM XIP (MGAS315 peptide)^19^ to induce natural competence and the ComRS pathway^16^ or 200 ng/mL of mitomycin C, to induce the phage to leave the genome (lytic cycle)^9^. XIP induced cells were grown for an additional 2 hours to an OD_600nm_ of 0.8. The mitomycin C induced samples decreased in optical density due to lysis. These cells were allowed to grow for an additional 4 hours after induction to an OD_600nm_ ranging from 0.2 to 0.8. The XIP triplicates and the mitomycin C induced triplicates were combined and cells harvested at 2000 *g*, 4 °C, for 20 min. The media was collected and cells re-suspended in 600 µL lysis buffer (50 mM Tris pH 7.5, 150 mM NaCl, 10% glycerol). The XIP and mitomycin C treated cells and divided evenly into 3 centrifuge tubes each (6 samples in total). The collected mitomycin C media was filtered with a 0.2 µm syringe filter and concentrated using a 3 kDa centrifugal filter (Amicon) to a volume of 500 µL. The cells and media were then flash-frozen and stored at -80 °C until use.

Frozen samples were thawed, 1 mM PMSF was added to each, and samples lysed by sonication using a (Qsonica XL-2000). Each GAS cell sample was approximately 250 µL for efficient sonication and the mitomycin C media sample was 500 µL. The cell samples were lysed by 3 rounds of sonication on ice. Each round was 1s on and 2s off with 30 second breaks. The media was lysed by one round to disrupt any phage particles. Lysed samples were then centrifuged at 21000 *g* for 10 min, at 4 °C, to remove insoluble material.

### Pull-down experiments of Prx with GAS lysates

To perform the pull-down experiments, each replicate of the XIP, mitomycin C, and mitomycin C media samples were subdivided into 2 centrifuge tubes of equal volume. To one tube 120 µg of purified Prx was added (+Prx), and to the second tube an equivalent volume of wash buffer (50 mM Tris pH 7.5, 150 mM NaCl, 20 mM imidazole, 10% glycerol) was added for the control (-Prx). Additionally, all sample volumes were increased to 700 µL with wash buffer to allow for more efficient sample mixing. Samples were allowed to incubate for 1.5 h at 4 °C while rotating. After incubation, 40 µL of Ni-NTA resin (Goldbio) prepared in wash buffer as a 50% slurry was added to each sample to pull-down Prx and any interaction partner. This provided a total bead volume of 20 µL. The samples were then allowed to incubate for an additional hour while mixing by rotation at 4°C. After incubation, beads were gently centrifuged at 1200x*g* for 5 min and the lysate removed. The beads were then washed 3 times with 600 µL wash buffer including mixing for 10 min at 4 °C. Each step was separated by centrifugation at 1200x*g* for 5 min to remove spent buffer. Finally, proteins were eluted with 40 µL of elution buffer (50 mM Tris pH 7.5, 150 mM NaCl, 500 mM imidazole, 10% glycerol) and beads removed by centrifugation at 1200x*g* for 10 min. The sample supernatants were carefully transferred to a new centrifuge tube, flash frozen, and stored at -80 °C.

### Protein digestion of GAS proteins for mass-spectrometry

A final concentration of 100 mM ammonium bicarbonate was added to each pull-down sample containing eluted peptides in total volume of 100 µL. Next peptides were reduced, by adding 10 µL of fresh 100mM DTT prepared in 100 mM ammonium bicarbonate. Samples were gently vortexed to mix, and allowed to incubate for 35 min at 58°C. After cooling to room temperature 10 µL 500 mM iodoacetamide (IAA) (Sigma) in 100 mM ammonium bicarbonate was added to alkylate the proteins. Samples were incubated for 45 min at room temperature protected from light. Next, 16 µL of 500 mM IAA was again added and allowed to react for 10 min to quench the excess IAA. Finally, sequencing grade trypsin (Promega) was added to each sample (2 µg). The samples were incubated overnight for 18h at 37°C to allow for complete protein digestion. The digested peptides were acidified and cleaned up for mass spectrometry using Pierce C18 spin columns (ThermoFisher: 89870) and the provided protocol. Briefly, the samples were acidified by adding 50 µL 4x stock sample buffer (2% Tri-fluoroacetic acid (TFA), 20% acetonitrile (ACN)), to 150 µL of each sample. The spin columns were prepared by washing with activation solution (50% ACN) 2 times followed by two washes with equilibration solution (0.5% TFA, 5% ACN). Samples were loaded and bound to the columns by centrifugation at 1500 *g* for 1 min. The flow-through was also re-run over the column. Peptide containing columns were washed 3 times with 200 µL equilibration solution and eluted with 20 µL of elution buffer (0.1% TFA, 70% ACN). The peptide samples were then stored at -20 °C before being analyzed by mass-spectrometry.

### Liquid chromatography with tandem mass-spectrometry of pull-down samples

Analysis of digests was carried out on an Orbitrap Exploris 480 instrument (Thermo Fisher Scientific, Bremen, Germany). The sample was introduced using an Easy-nLC 1000 system (Thermo Fisher Scientific) at 830 ng per injection. Mobile phase A was 0.1% (v/v) formic acid and mobile phase B was 0.1% (v/v) formic acid in 80% acetonitrile (LC-MS grade). Gradient separation of peptides was performed on a C18 [Luna C18(2), 3 μm particle size (Phenomenex, Torrance, CA)] column packed in-house in Pico-Frit (100 μm X 30 cm) capillaries (New Objective, Woburn, MA). Peptide separation was conducted using the following gradients: started with 2% of phase B, 2 – 6% over 5 minutes, 6 – 25% over 64 minutes, 25 – 45% over 10 minutes, 45 – 90% over 1 minute, with final elution of 90% B for 15 minutes at a flow rate of 300 nL/minute.

Data acquisition on the Orbitrap Exploris 480 instrument was configured for data-dependent method using the full MS/DD−MS/MS setup in a positive mode. Spray voltage was set to 2.7 kV, funnel RF level at 40, and heated capillary at 275°C. Survey scans covering the mass range of 380–1500 m/z were acquired at a resolution of 60,000 (at m/z 200), with a normalized automatic gain control (AGC) target of 300% and an auto maximum ion injection time. This was followed by MS2 acquisition at a resolution of 15,000 with an intensity threshold kept at 2e4. During MS2 acquisition, the 20 most abundant ions were selected for fragmentation at 30% normalized collision energy. AGC target value for fragment spectra was set to standard with a maximum ion injection time set to auto and an isolation width set at 1.6 m/z. Dynamic exclusion of previously selected masses was enabled for 20 seconds, charge state filtering was limited to 2–6, peptide match was set to preferred, and isotope exclusion was on.

### Mass-spectrometry data analysis and protein identification

Raw spectra files were converted into Mascot Generic File format (MGF) for peptide/protein identification by X!Tandem search algorithm^52^. The following X!Tandem search parameters were used: 10 ppm and 20 ppm mass tolerance for parent and fragment ions, respectively; constant modification of Cys with iodoacetamide; default set of variable post-translational modifications: oxidation of Met, Trp; N-terminal cyclization at Gln, Cys; N-terminal acetylation, phosphorylation (Ser, Thr, Tyr), deamidation (Asn and Gln); an expectation value cut-off of Log(e) < –1 for peptides was used, corresponding to ∼0.3% false discovery rate. Protein identification cut-off was set for Log(e) < –8 and at least 2 peptides per protein.

### Size exclusion chromatography binding assays

SEC protein binding assays were performed using an AKTApure (Cytiva), using either a Superdex75 increase 10/300 column (Cytiva) for EndoS1, Prx, JM3, and EndoS1 and Prx variants, or a Superdex200 increase 10/300 column (Cytiva) for EndoS1, HsdS and HsdM. All assays were performed at 4°C. The proteins for each assay were incubated together or alone in assay buffer (50 mM HEPES at pH 7.5, 150 mM NaCl, I mM β-Me) on ice for 30 min before loading onto the appropriate SEC column equilibrated in assay buffer. Protein concentrations were set to a molar ratio of 2:1 for small molecular weight to high molecular weight to allow for clear separation of earlier eluting complex peaks. After each run, fractions were collected and analysed by Coomassie stained SDS-PAGE gel.

### Isothermal titration calorimetry

EndoS1, Prx, and Prx residue variants were dialyzed overnight into the same buffer (50 mM HEPES at pH 7.5, 150 mM NaCl, I mM β-Me). ITC was performed using a MicroCal ITC200 (Malvern). Each experiment required ∼40 uL of 600 µM of Prx injected to ∼240 uL of 50 µM EndoS1 at a constant temperature of 25°C. Controls were run with buffer injected into EndoS1 and Prx injected into buffer. Data was evaluated using the Origin Software (Malvern) and binding constants and thermodynamic values calculated using a one-site model.

### Protein crystallization

Crystallization conditions were screened using commercially available crystal screens (NeXtal, Molecular Dimensions) with a Crystal Gryphon Robot (Art Robbins Instruments) or manually using a multichannel pipette. For both purified EndoS1 and an N-terminal truncated EndoS1:Prx complex, conditions were screened at a protein concentration of 12 mg/mL, 8 mg/mL and 4 mg/mL. Proteins were mixed with mother liquor at a 1:1 ratio, setup using the sitting drop vapor diffusion method and incubated at 4°C. EndoS1 crystals were obtained in 0.2 M Lithium sulfate, 0.1 M Tris pH 8.5, 40% PEG 200. EndoS1:Prx cocrystals were obtained in 0.2 M sodium chloride, 0.1M sodium phosphate pH 6.2, 50% PEG 200. Crystals were found to grow at all tested protein concentrations.

### Data collection and refinement

The data set for EndoS1 was collected at the Canadian Light Source beamline CMCF-BM (08B1). EndoS1 crystals did not require cryo-protection. The data set for EndoS1:Prx was collected in house at 93K using a MicroMax-007 HF X-ray source and R-axis 4++ detector (Rigaku). EndoS1:Prx crystals did not require additional cryo-protection. All protein crystals were flash frozen directly in liquid nitrogen before mounting. Data was processed using XDS^53^ and CCP4^54^. Initial phases for EndoS1 were obtained with Phenix^55^ by molecular replacement utilizing a model of EndoS1 predicted by AlphaFold^33^. Phases for the EndoS1:Prx complex were obtained my molecular replacement using the EndoS1 model and Prx (PDBid 6CKA). Proteins models were built in Coot and refined with Phenix^55^, Refmac5^56^, and TLS^57^. ChimeraX^58^ was utilized for molecular graphics.

## DATA AVAILABILITY

The X-ray structure and diffraction data reported in this paper have been deposited in the Protein Data Bank under the accession codes 8VSQ for EndoS1 and 8VSR for EndoS1:Prx. Mass-spectrometry raw data has been deposited in the University of San Diego MassIVE site (massive.ucsd.edu) with accession number MSV00009549.

## SUPPORTING INFORMATION

This article contains supporting information.

## ACKNOWLEDGEMENTS

We would like to thank the laboratory of Jörg Stetefeld and Markus Meier at the University of Manitoba for instrument access and assistance with SEC-MALS. Additionally, we thank René Zahedi and Victor Spicer at the University of Manitoba for assistance in mass-spectrometry data interpretation and deposition. We would also like to thank beamline CMCF-BM at the Canadian Light Source, which is supported by the Canada Foundation for Innovation (CFI), the Natural Sciences and Engineering Research Council (NSERC), the National Research Council (NRC), the Canadian Institutes of Health Research (CIHR), the Government of Saskatchewan.

## FUNDING AND ADDITIONAL INFORMATION

This work was supported by the Natural Sciences and Engineering Research Council of Canada (NSERC) Discovery grant RGPIN-2018-04968 to G.P., a Manitoba Medical Service Foundation Operating Grant (MMSF) 8-2022-10 to G.P., and a Canadian Foundation for Innovation award (CFI) 37841 to G.P. This work was also supported by a University of Manitoba Research Grants Program (URGP) to GP.

