## Supplemental_data for "The phage protein paratox is a multifunctional metabolic regulator of *Streptococcus*"

Running title: Paratox binds multiple transcriptional regulators in Streptococcus

**Key words:**
Paratox, Prx, ComR, *Streptococcus,* quorum sensing, bacteriophage, X-ray crystallography, mass-spectrometry, transcription

* To whom correspondence should be addressed: G.P.


List of Supplementary information:

Table S1
Figures S1-S11

**Table S1: Primers used in this study**

| | |
|---|---|
| P140 | ACGTCATATGGAAAATAGCATTGAGGCCTATCTAG |
| P141 | CGTACTCGAGACCTAATCCTTTTTGAATTTCATTCTGAATATATTC |
| P152 | AGGTACCATATGACAAAATCTAAACAGCCTCAATATCGC |
| P154 | CATATGTATATCTCCTTCTTAAAGTTAAACAAA ATTATTTC |
| P155 | GCAGATAGCAGAGAAAAGGTAACATTAG |
| P158 | TCCTTTTTGAATTTCATTCTGAATATATTCCCACTCCTG |
| P159 | AGGTACCATATGGCAGAGAAAACAACTTCACTCC |
| P183 | TACAGCGGCCGCTTAGTTTCTAAAAGTATCCAATTCGTCTTG |
| P202 | GCTAAGGATCCATGAAAAGACACAGACAGTTTAATAAAGATATTAAATAC |
| P203 | CGATTGCGGCCGCTCAGCATCTCTCCACCTCTCTC |
| P206 | CGTACTCGAGTTATACGAATAAACGATTGAGTAATGTTTG |
| P207 | ATCTGAATTCGATGGCAGAGAAAACAACTTCAC |
| P211 | CCTACGATCCATCTGGAAAGTTC |
| P212 | GCGCTCTTGCGTTACCTTTTGGAAG |
| P213 | CAAGAGCTGCCTACGATCCATC |
| P214 | GCTTACCTTTTGGAAGACGGAGATGTC |

| Protein | Uniprot identifier | -Prx | + Prx | Annotation |
|---|---|---|---|---|
| SpyM3_0271 | A0A0H2UTH1 | / | -108.9 | putative tRNA (cytidine(34)-2'-O)-methyltransferase |
| SpyM3_0575 | P0DB06 | / | -107.2 | NAD(P)/FAD-dependent oxidoreductase |
| SpyM3_1203 | A0A0H2UVL5 | / | -91 | Paratox Spy_1203 (315.4) |
| SpyM3_1408 | A0A0H2UW47 | / | -85.8 | Paratox Spy_1408 (315.6) |
| SpyM3_0890 | A0A0H2UUU0 | / | -81.6 | type I restriction endonuclease subunit S/type I restriction endonuclease subunit S |
| SpyM3_0977 | A0A0H2UVP7 | / | -63.5 | LexA family transcriptional regulator/putative repressor protein - phage-associated |
| SpyM3_0500 | P0CZ96 | -1.7 | -58.5 | ATP synthase F1, epsilon subunit |
| SpyM3_0191 | P0DF42 | / | -53 | ribosome-associated GTPase .YjeQ_engC, YjeQ/EngC. |
| SpyM3_0246 | P0DC74 | / | -40.1 | Transcriptional repressor NrdR |
| SpyM3_0843 | A0A0H2UUF4 | / | -36 | GntR family transcriptional regulator/ UbiC transcription regulator-associated domain protein |
| SpyM3_1036 | A0A0H2UVU0 | -2.5 | -26.4 | biotin-(acetyl-CoA-carboxylase) ligase / BirA |
| SpyM3_0573 | P0DF04 | / | -20.4 | Ribosome maturation factor RimM/ 16s rRNA processing protein |
| SpyM3_1027 | P0CZ72 | / | -19 | 3-phosphoshikimate 1-carboxyvinyltransferase |
| SpyM3_1619 | P0DH18 | / | -17.9 | Putative transcriptional regulator/DNA-dependent RNA polymerase auxiliary subunit epsilon family protein |
| SpyM3_0418 | A0A0H2UU97 | / | -16.5 | U32 family peptidase / collagenase-like protease |
| E0F66_03415 | A0A5S4TSJ5 | / | -16.2 | tRNA O-methyltransferase TrmR |
| SpyM3_0318 | P0DF60 | -1.9 | -13.1 | putative metal binding protein of ABC transporter/manganese-binding protein |
| SpyM3_0888 | A0A0H2UUI4 | / | -11.8 | Duf2130 domain containing protein/ hypothetical prot SpyM3_0888 |
| E0F66_00310 | A7XFU9 | / | -10.9 | hylA:p, extracellular hyaluronate lyase .GAG_Lyase, Glycosaminoglycan (GAG) polysaccharide lyase |
| SpyM3_0307 | A0A0H2UU08 | / | -10.9 | hypothetical phage protein/ SpyM3_0307 |
| SpyM3_1470 | P0DG98 | -2.1 | -10.3 | hypothetical prot SpyM3_1470 / DUF896 |
| SpyM3_0455 | A0A0H2UTT1 | / | -10 | ftsE:p, putative cell-division ATP-binding protein |
| SpyM3_1837 | A0A0H2UXZ4 | / | -8.4 | DHH family phosphoesterase / SpyM3_1837 |

**Figure S1: Prx binding partners identified from *Streptococcus pyogenes* stimulated with XIP.** Purified Prx bound to nickel-agarose beads (+Prx) was used to co-precipitate proteins from GAS extracts stimulated with XIP (GAS natural competence). Empty beads (-Prx) was used as a control. Protein hits were identified by mass-spectrometry and are listed by protein name, Uniprot identifier, and putative annotations. Data is reported as the Log expectation value, where a larger negative value indicates a higher confidence protein hit. No peptides detected by mass-spectrometry is indicated by a backslash.

| Protein | Uniprot identifier | -Prx | +Prx | Annotation |
|---|---|---|---|---|
| SpyM3_0575 | P0DB06 | / | -207.6 | NAD(P)/FAD-dependent oxidoreductase/ thioredoxin reductase |
| SpyM3_0191 | P0DF42 | / | -135.2 | ribosome-associated GTPase .YjeQ_engC, YjeQ/EngC. YjeQ (YloQ in Bacillus subtilis) |
| SpyM3_0573 | P0DF04 | / | -107.8 | ribosome maturation factor RimM / 16s rRNA processing protien |
| SpyM3_0890 | A0A0H2UUU0 | / | -85.3 | type I restriction endonuclease subunit S / SpyM3_0890 |
| SpyM3_1203 | A0A0H2UVL5 | -1.9 | -81.1 | Paratox SpyM3_1203 (315.4) |
| SpyM3_0843 | A0A0H2UUF4 | / | -73.9 | GntR family transcriptional regulator/ UbiC transcription regulator-associated domain protein |
| SpyM3_0810 | P0DG02 | / | -65.5 | gyrA:p, DNA gyrase subunit A .PRK05560, DNA gyrase subunit ADNA_gyraseA_C |
| SpyM3_0634 | P0DC32 | / | -61.1 | tRNA (adenosine(37)-N6)-dimethylallyltransferase MiaA |
| SpyM3_1408 | A0A0H2UW47 | / | -56.9 | Paratox SpyM3_1408 (315.6) |
| SpyM3_0307 | A0A0H2UU08 | / | -47 | hypothetical protein SpyM3_0307 / phage protein |
| E0F66_03415 | A0A5S4TSJ5 | / | -45.1 | tRNA O-methyltransferase TrmR |
| SpyM3_1499 | P0DF06 | / | -44.6 | ribosome maturation factor RimP |
| SpyM3_0635 | A0A0H2UTY2 | / | -37.4 | GTP-binding protein HflX |
| SpyM3_1030 | A0A0H2UUU3 | / | -32.1 | Cytosolic protein containing multiple CBS domains /transcription factor SpxR |
| SpyM3_0737 | P0DC22 | -2 | -27.8 | lepA:p, GTP-binding protein LepA .lepA_C, lepA_C |
| SpyM3_0964/1346 | A0A0H2UUS6 | / | -25.8 | XRE family transcriptional regulator / cro/cI family (phage, SpyM3_0964, SpyM3_1346) |
| SpyM3_1036 | A0A0H2UVU0 | / | -25.7 | birA:p, biotin--protein ligase syntheta...BPL_C, Biotin protein ligase C terminal domain |
| SpyM3_1246 | A0A0H2UVB1 | -1.9 | -23.1 | hypothetical phage protein SpyM3_1246 |
| SpyM3_0031 | A0A0H2USW3 | / | -22.5 | helix-turn-helix transcriptional regulator/XRE family |
| SpyM3_0539 | P0DE48 | / | -17.7 | rpmI:p, 50S ribosomal protein L35 .Ribosomal_L35p, Ribosomal protein L35 |
| SpyM3_1248 | A0A0H2UVP5 | / | -16.8 | RusA crossover junction endodeoxyribonuclease/ holiday junction resolvase/ phage protein |
| SpyM3_0418 | A0A0H2UU97 | -1.5 | -16.5 | U32 family peptidase / collagenase-like protease |
| SpyM3_1618 | A0A0H2UXH6 | / | -15.8 | putative glycoprotein endopeptidase .COG1214 |
| E0F66_02980 | A0A5S4TMV3 | -2.2 | -15 | hypothetical protein SpyM3_1149 .DegV, Uncharacterised protein, DegV family COG1307 |
| SpyM3_0476 | P0DG04 | -1.2 | -14.5 | gyrB:p, DNA gyrase subunit B |
| SpyM3_1094 | A0A0H2UVY3 | / | -14.4 | Paratox SpyM3_1094 (315.3) |
| SpyM3_1837 | A0A0H2UXZ4 | / | -12.2 | DHH family phosphoesterase |
| SpyM3_0434 | A0A0H2UTL9 | -1.2 | -12.1 | aminoacyltransferase/ femAB family protein |
| SpyM3_1800 | P0DD82 | -2.2 | -10.7 | recombinaseA (RecA) |
| SpyM3_0364 | A0A0H2UTG1 | / | -10.6 | 1,2-diacylglycerol 3-glucosyltransferase |
| SpyM3_0153 | P0DA20 | / | -10.5 | tRNA adenosine(34) deaminase TadA |
| SpyM3_0229 | A0A0H2UT77 | / | -10.2 | class I SAM-dependent methyltransferase |
| SpyM3_0588 | A0A0H2UU36 | / | -10 | ABC transporter ATP-binding protein |
| SpyM3_0691 | A0A0H2UU60 | / | -9.8 | hypothetical phage protein SpyM3_0691 / replication initiator A protien |
| SpyM3_0888 | A0A0H2UUI4 | / | -9.1 | DUF2130 domain-containing protein / hypotheical SpyM3_0888 |
| SpyM3_1470 | P0DG98 | / | -8.5 | Bacterial protein of unknown function (DUF896) |
| SpyM3_1858 | P0DG60 | / | -8.3 | trsA:p, tryptophanyl-tRNA synthetase II |

**Figure S2: Prx binding partners identified from *Streptococcus pyogenes* stimulated with mitomycin C.** Purified Prx bound to nickel-agarose beads (+Prx) was used to co-precipitate proteins from GAS extracts stimulated with mitomycin C to induce phage protein expression and lytic exit. Empty beads (-Prx) was used as a control. Protein hits were identified by mass-spectrometry and are listed by protein name, Uniprot identifier, and putative annotations. Data is reported as the Log expectation value, where a larger negative value indicates a higher confidence protein hit. No peptides detected by mass-spectrometry is indicated by a backslash.

| Protein | Uniprot Identifier | -Prx | + Prx | Annotation |
|---|---|---|---|---|
| SpyM3_0575 | P0DB06 | / | -184 | Ferredoxin--NADP reductase |
| SpyM3_0890 | A0A0H2UUU0 | / | -147 | Uncharacterized protein, S-subunit endonuclease |
| SpyM3_0455 | A0A0H2UTT1 | / | -144 | Cell division ATP-binding protein FtsE |
| SpyM3_0810 | P0DG02 | / | -136 | DNA gyrase subunit A |
| SpyM3_0271 | A0A0H2UTH1 | / | -135 | Putative tRNA (cytidine(34)-2'-O)-methyltransferase |
| SpyM3_1433 | A0A0H2UW63 | / | -128 | Phage head morphogenesis domain-containing protein |
| SpyM3_0635 | A0A0H2UTY2 | / | -125 | GTPase HflX |
| SpyM3_1588 | A0A0H2UWA4 | / | -91 | Endonuclease MutS2 |
| SpyM3_0573 | P0DF04 | / | -87 | Ribosome maturation factor RimM |
| SpyM3_0364 | A0A0H2UTG1 | / | -86.7 | Putative sugar transferase |
| SpyM3_0191 | P0DF42 | / | -85.7 | Small ribosomal subunit biogenesis GTPase RsgA |
| SpyM3_0461 | P0DG28 | / | -83.9 | Asparagine--tRNA ligase |
| SpyM3_1203 | A0A0H2UVL5 | / | -83.4 | Paratox |
| SpyM3_1060 | A0A0H2UUW8 | / | -80.9 | tRNA 5-hydroxyuridine methyltransferase |
| SpyM3_0516 | P0DE86 | / | -75.8 | 30S ribosomal protein bS21 |
| SpyM3_0964 | A0A0H2UUS6 | / | -74.9 | HTH cro/C1-type domain-containing protein |
| SpyM3_0977 | A0A0H2UVP7 | / | -66.1 | Putative repressor protein-phage-associated |
| SpyM3_0500 | P0CZ96 | / | -64.3 | ATP synthase epsilon chain |
| SpyM3_0634 | P0DC32 | / | -58.4 | tRNA dimethylallyltransferase |
| SpyM3_1499 | P0DF06 | / | -58 | Ribosome maturation factor RimP |
| SpyM3_1633 | P0DF34 | / | -55.4 | Probable DNA-directed RNA polymerase subunit delta |
| SpyM3_0749 | P0DH40 | / | -55.2 | UvrABC system protein C |
| SpyM3_1030 | A0A0H2UUU3 | / | -49.3 | CBS domain-containing protein |
| SpyM3_0246 | P0DC74 | / | -48.9 | Transcriptional repressor NrdR |
| SpyM3_1036 | A0A0H2UVU0 | / | -41 | Bifunctional ligase/repressor BirA |
| SpyM3_1211 | A0A0H2UV87 | / | -37.3 | Phage protein |
| SpyM3_0049 | P0DE78 | / | -34.9 | 30S ribosomal protein S17 |
| SpyM3_0539 | P0DE48 | / | -34.3 | 50S ribosomal protein L35 |
| SpyM3_1374 | P0DF36 | / | -33 | DNA-directed RNA polymerase subunit omega |
| SpyM3_1619 | P0DH18 | / | -31.6 | DNA-directed RNA polymerase subunit epsilon |
| SpyM3_1816 | P0DE40 | / | -31.3 | 50S ribosomal protein L32 |
| SpyM3_1389 | P0DD90 | / | -31 | Holliday junction resolvase RecU |
| SpyM3_1629 | P0DE30 | / | -26.9 | 50S ribosomal protein L28 |
| SpyM3_1246 | A0A0H2UVB1 | / | -26.5 | Phage protein |
| SpyM3_0665 | A0A0H2UU33 | / | -24.9 | Extracellular hyaluronate lyase |
| SpyM3_0657 | Q8K7S6 | / | -24.1 | Ribonuclease J 2 |
| SpyM3_1283 | A0A0H2UWM0 | / | -22.2 | Putative methyltransferase |
| SpyM3_0478 | A0A0H2UTQ0 | / | -19.3 | DUF1694 domain-containing protein |
| SpyM3_1580 | P0DE80 | / | -17.3 | 30S ribosomal protein S18 |
| SpyM3_0051 | P0DE26 | / | -16.4 | 50S ribosomal protein L24 |
| SpyM3_1500 | P0DG18 | / | -16 | tRNA (guanine-N(7)-)-methyltransferase |
| SpyM3_0169 | A0A0H2UT49 | / | -14.9 | Uncharacterized protein |
| SpyM3_0196 | A0A0H2UT59 | / | -13.2 | Pur operon repressor |

**Figure S3: Prx binding partners identified from *Streptococcus pyogenes* growth media after cells were stimulated with mitomycin C.** Purified Prx bound to nickel-agarose beads (+Prx) was used to co-precipitate proteins found in the lysate of GAS cells that had been stimulated with mitomycin C to induce phage protein expression and lytic exit. Empty beads (-Prx) was used as a control. Protein hits were identified by mass-spectrometry and are listed by protein name, Uniprot identifier, and putative annotations. Data is reported as the Log expectation value, where a larger negative value indicates a higher confidence protein hit. No peptides detected by mass-spectrometry is indicated by a backslash.

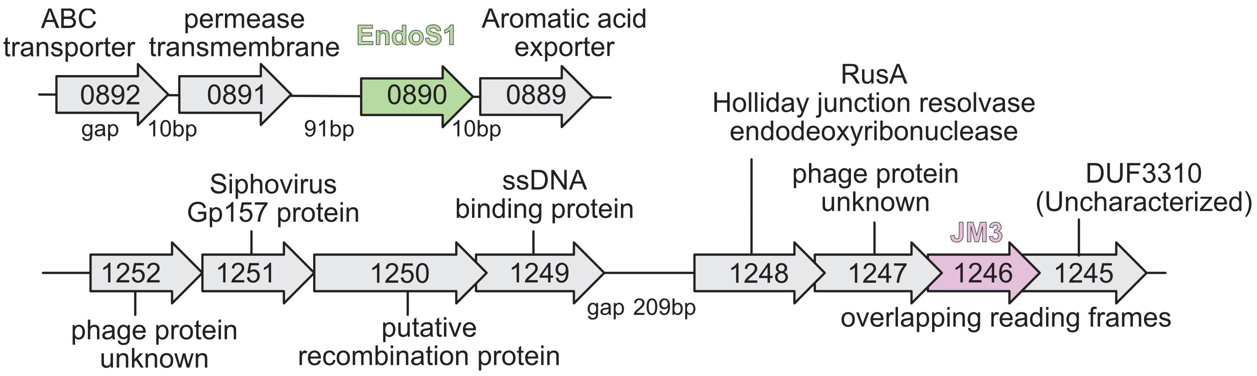


**Figure S4: Map of the local genomic regions for the genes encoding Prx binding partners EndoS1 and JM3.** (Top): The gene encoding EndoS1 (*spyM3_0890*) is located in the GAS chromosome near several genes that are annotated to encode metabolite transporters and membrane permeases. (Bottom): The gene encoding JM3 (*spyM3_1246*) is located within the prophage and is part of a cluster of genes predicted to be involved in DNA recombination and resistance to other phages (Gp157 protein).

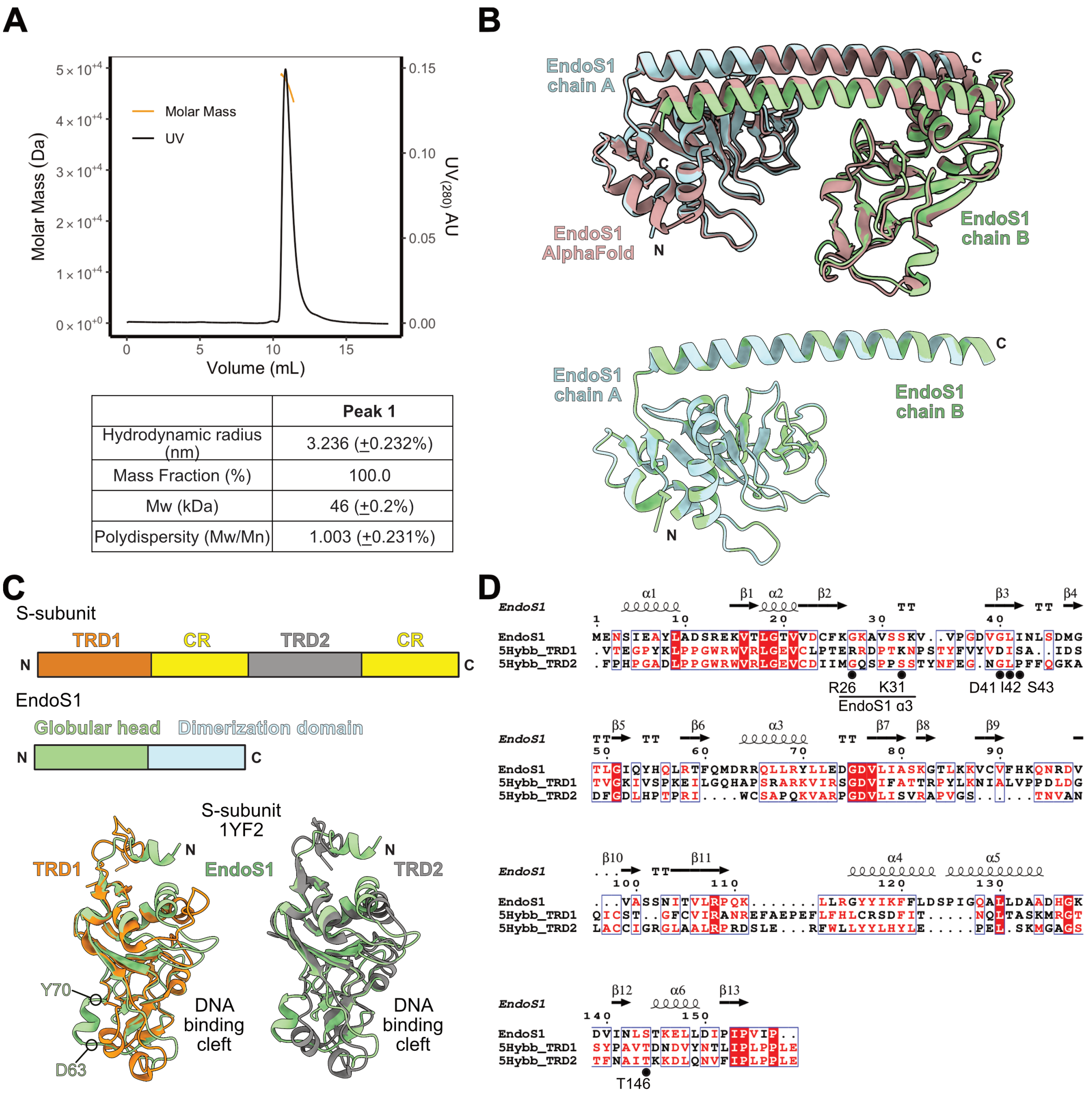


| | Peak 1 |
|---|---|
| Hydrodynamic radius (nm) | 3.236 (±0.232%) |
| Mass Fraction (%) | 100.0 |
| Mw (kDa) | 46 (±0.2%) |
| Polydispersity (Mw/Mn) | 1.003 (±0.231%) |

**Figure S5: Structural analysis of EndoS1. (A)** EndoS1 is a dimer in solution as assayed by SEC-MALS. ASTRA software (Wyatt Technologies) analysis shows a single species in solution of 46 kDa, or twice the theoretical molecular weight of the 23 kDa EndoS1 monomer. **(B)** Top: Structural alignment of the monomeric AlphaFold model for EndoS1(brown) with each chain in the experimental EndoS1 dimer structure. The AlphaFold model was provided by the Uniprot entry for EndoS1 (SpyM3_0890). Bottom: Structural alignment of each chain in from the EndoS1 crystal structure. **(C)** Structural comparison of EndoS1 with classical restriction endonuclease S-subunits. Top: Cartoon domain map of an S-subunit compared to EndoS1. TRDs are colored orange and grey, with the conserved regions (CR) in yellow. The EndoS1 globular head (green) corresponds to a TRD and the dimerization domain (blue) corresponds to a CR. Bottom: structural alignment of the EndoS1 globular head to the TRD domains of S-subunit PDBid 1YF2. **(D)** Sequence alignment of EndoS1 to the TRDs of S-subunit PDBid 5HYBB. Numbering is based on the sequence of EndoS1. Residues important for DNA binding in TRD1 are labelled and indicated by a black circle. R26 and K31 are found structurally aligned to the third alpha-helix of EndoS1 (EndoS1 α3 or insertion helix) and not apparent from the sequence alignment. Sequence alignment was generated by the server ClustalOmega and colored by conservation using the server Espript3 (https://espript.ibcp.fr/ESPript/ESPript/).

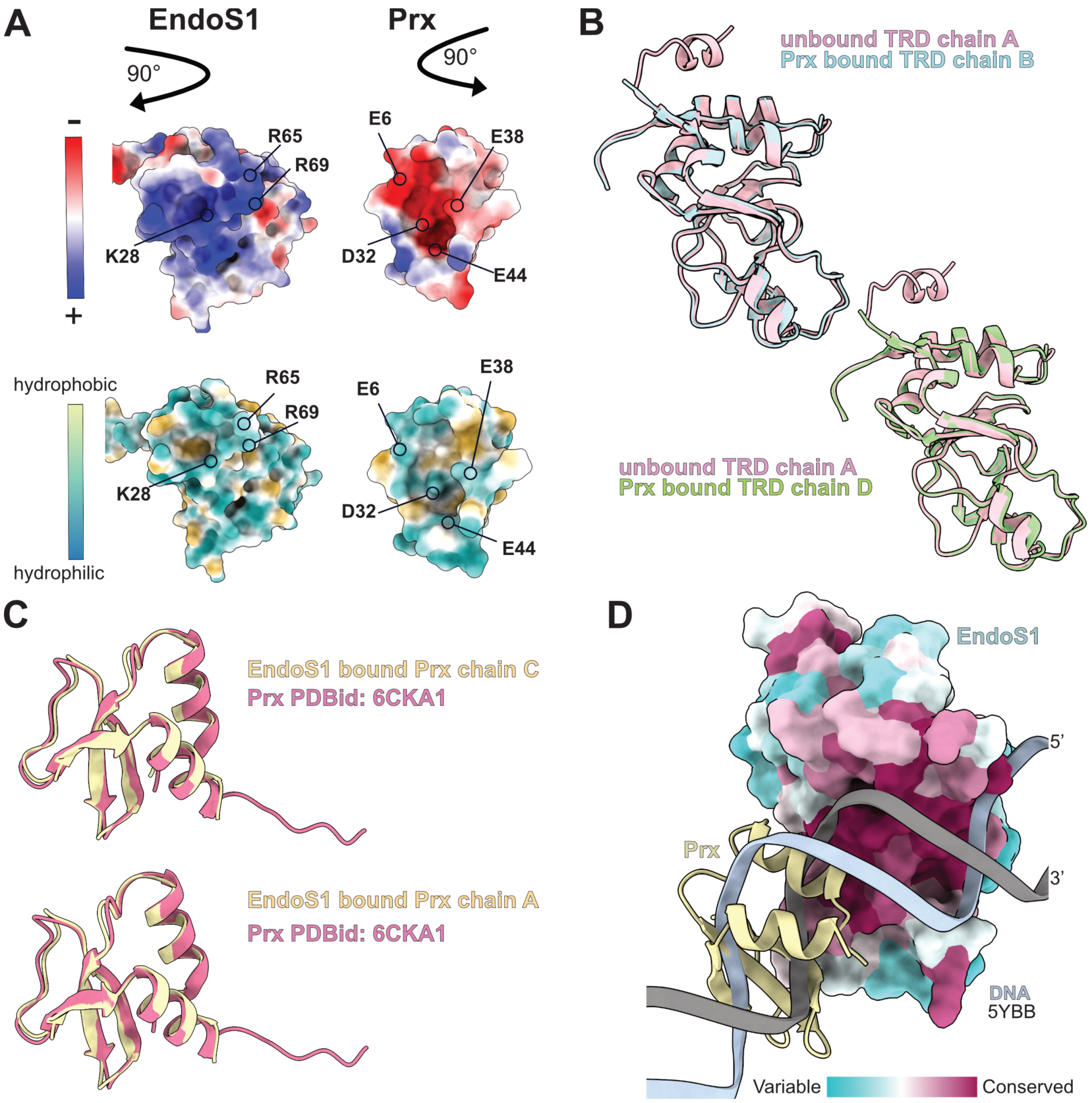


**Figure S6: Structural analysis of an EndoS1:Prx cocrystal complex. (A)** Molecular surface properties of the EndoS1:Prx interaction. Each panel shows the contact surface on each protein with important residues outlined in Figure 5B highlighted. Top: Relative electrostatics colored by charge showing complementary properties. Bottom: Hydrophobic surfaces on each protein demonstrating a primarily polar interface. **(B)** Overlay of the globular heads of unbound EndoS1 and EndoS1:Prx showing no local conformational changes in the protein fold. **(C)** Overlay of Prx (PDBid: 6CKA1) with Prx that is in complex with EndoS1 demonstrated the fold of Prx is identical. **(D)** Structural alignment of EndoS1:Prx with a DNA bound S-subunit from 5YBB as in Figure 4E. The alignment shows that Prx binding EndoS1 would prevent EndoS1 from binding DNA.

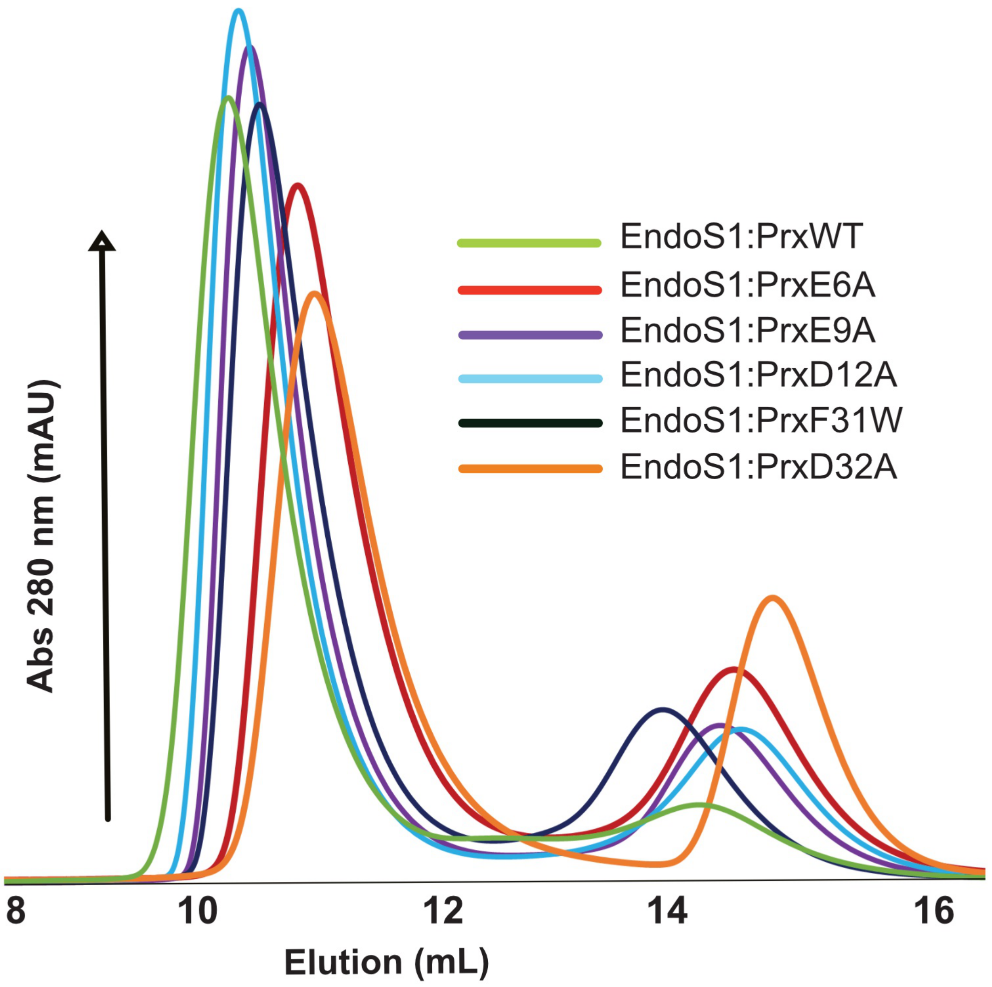


**Figure S7: Complete Prx variant binding assays with purified EndoS1.** SEC binding assay of EndoS1 with different Prx variants. EndoS1 was mixed with each Prx variant separately and loaded over a Superdex75 increase 10/300 GL SEC column (Cytiva). The expected shift in elution volume for complex formation with EndoS1 with wild type Prx is shown in green.

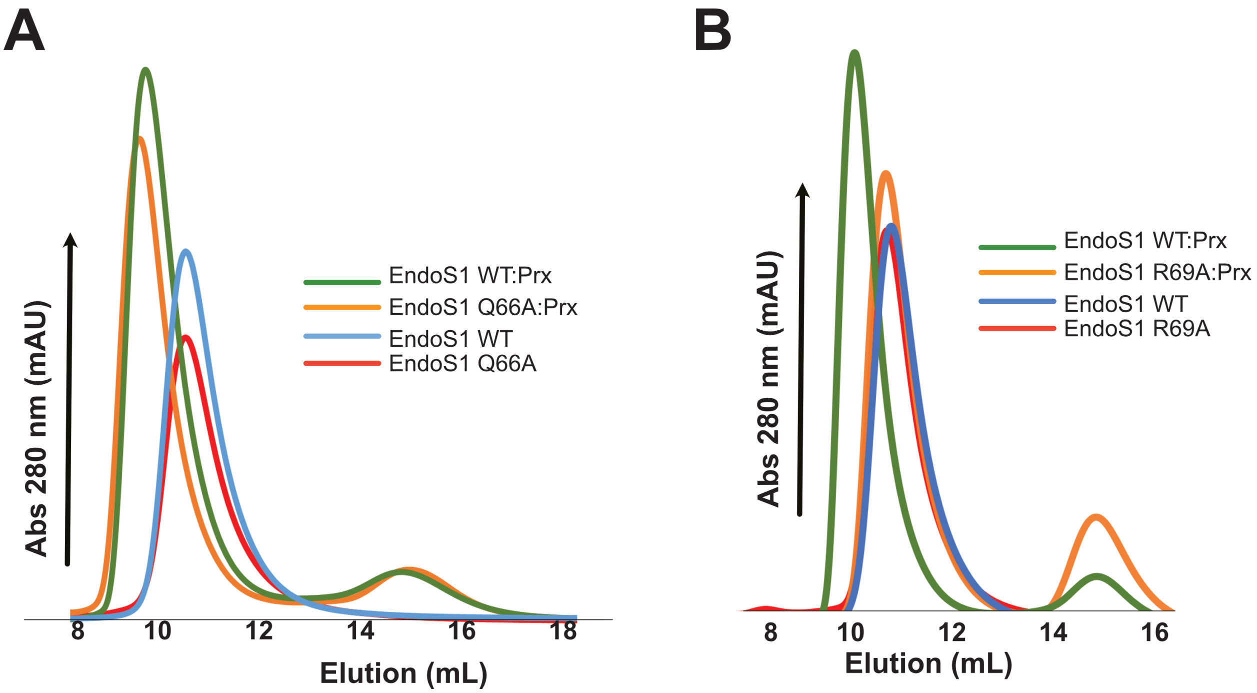


**Figure S8: SEC traces of EndoS1 variant binding assays with Prx. (A)** Overlay of chromatograms for EndoS1 (blue), EndoS1Q66A (red), EndoS1:Prx (green), and EndoS1Q66A:Prx (orange). **(B)** Overlay of chromatograms for EndoS1 (blue), EndoS1R69A (red), EndoS1:Prx (green), and EndoS1R69A:Prx (orange). All experiments were performed using a Superdex75 increase 10/300 GL SEC column (Cytiva).

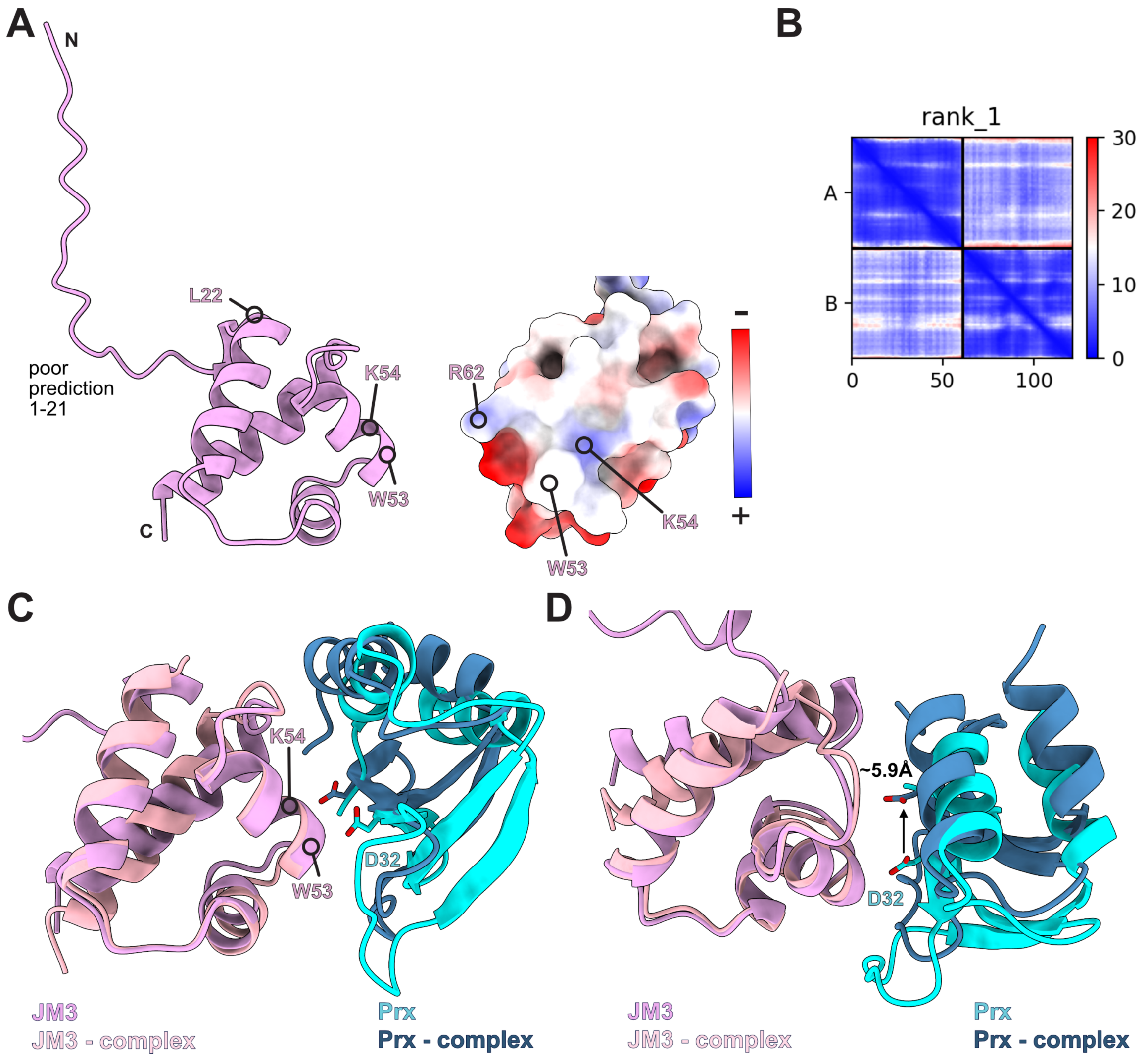


**Figure S9: AlphaFold models of JM3 and a JM3:Prx complex. (A)** AlphaFold model of JM3 available from Uniprot. The cartoon depiction (left) and relative electrostatics of the putative molecular surface interface with Prx are shown (right). L22 represents the start of the helix-turn-helix (HTH) DBD domain. Residues at the predicted Prx interface are also shown. The right image is a 90° rotation from the left image. **(B)** Predicted aligned error (PAE) plots for the top JM3:Prx complex predicted by AlphaFold2. The complex was predicted using the AlphaFold2 google colab. **(C)** Structural comparison of ComR:Prx (PDBid 7N10) aligned to JM3 (violet:cyan) and the JM3:Prx AlphaFold2 prediction (peach:dark blue). **(D)** Same structures as panel **(C)** but rotated 90° showing that AlphaFold predicts a similar but shifted binding interface relative to how Prx binds the ComR DBD.

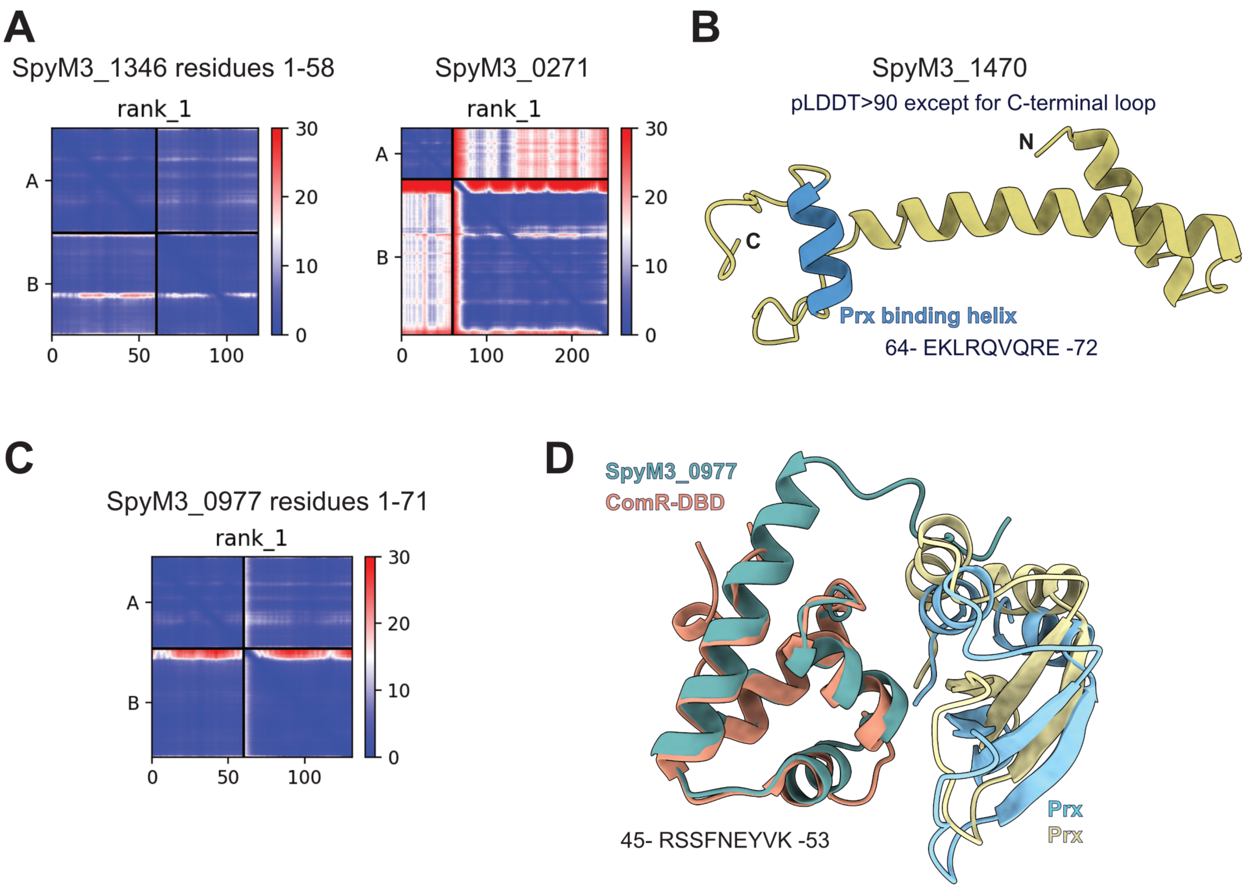


**Figure S10: AlphaFold models of Prx binding partners in complex with Prx. (A)** Predicted aligned error (PAE) plots for the top models of the SpyM3_1346:Prx complex and the SpyM3_0271:Prx complex. **(B)** AlphaFold model of SpyM3_1470 as provided by Uniprot. The putative Prx binding helix is colored in blue. The AlphaFold prediction is of high confidence (pLDDT>90) for the whole model except the C-terminal loop. **(C)** Predicted aligned error (PAE) plots for the top AlphaFold model of SpyM3_0977 with Prx. **(D)** ComR:Prx (PDBid 7N10) (orange:pale-goldenrod) structural aligned to a SpyM3_0977:Prx complex predicted by AlphaFold (aqua-green:cyan). SpyM3_0977 contains a helix-turn-helix domain fold nearly identical to ComR, JM3, and SpyM3_1346. SpyM3_1346 lacks the Prx binding motif (Figure 8) similar to JM3. However, like JM3 AlphaFold predicts a similar but modified Prx binding surface.

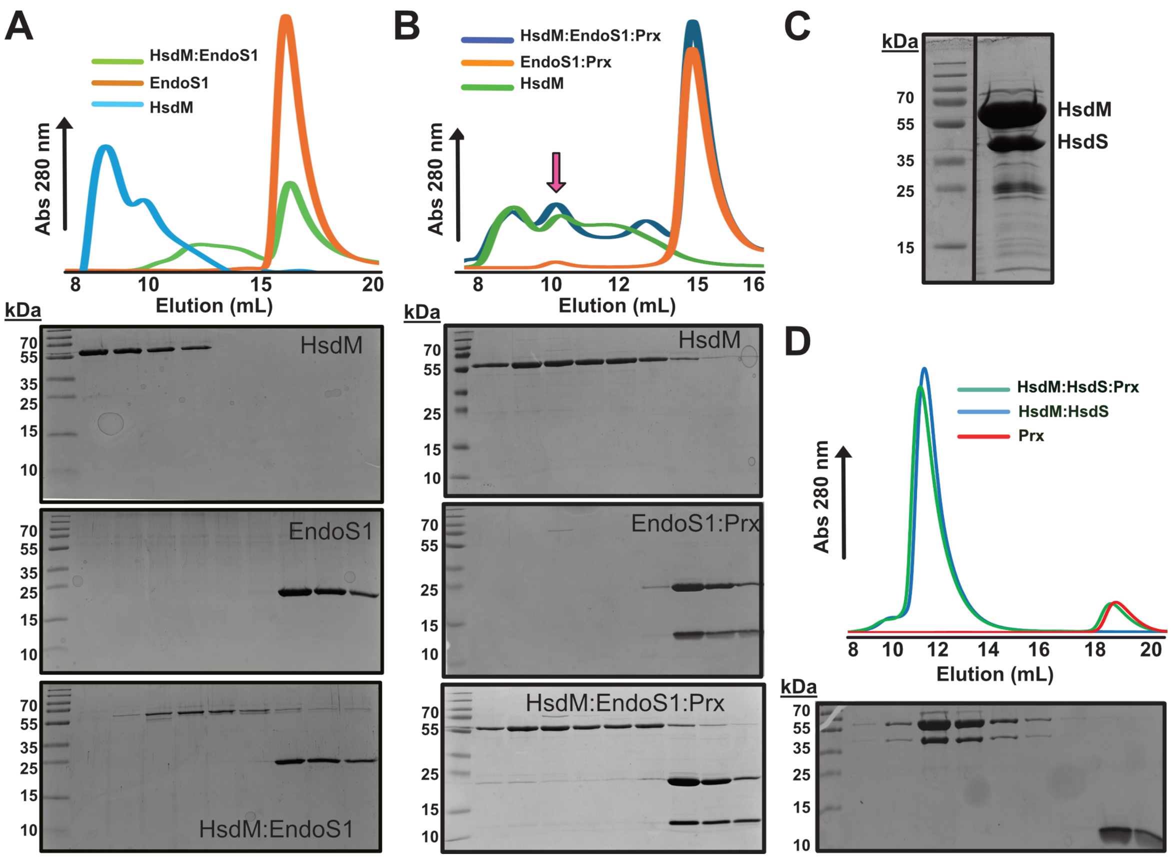


**Figure S11: SEC binding assays of EndoS1 and Prx with HsdS and HsdM. (A)** Binding experiment between EndoS1 and HsdM. The top panel shows SEC traces of EndoS1 (orange), HsdM (blue), and HsdM mixed with EndoS1 (green). The bottom panel shows Coomassie-stained SDS-PAGE gels of each experiment. The trace is aligned to the gels based on elution volume. The behavior of HsdM changes in the presence of EndoS1 but no shift and stable complex is observed. **(B)** Binding experiment between HsdM (green) and an EndoS1:Prx complex (orange). Top and bottom panels are as described in (A). The magenta arrow highlights a potential HsdM:EndoS1:Prx complex (blue) as the SDS-PAGE gel seems to suggest a shift of the EndoS1:Prx complex. However, this result was not reliably reproducible. **(C)** Purified HsdM:HsdS co-expression complex as purified by metal affinity chromatography. **(D)** Binding experiment between an HsdM:HsdS complex (blue) and Prx (red). The HsdM:HsdS complex is a single stable peak that does not interact with Prx (green). Top and bottom panels are also as described in (A).
